# Structural and Biophysical Dynamics of Fungal Plasma Membrane Proteins and Implications for Echinocandin Action in *Candida glabrata*

**DOI:** 10.1101/2024.05.29.596243

**Authors:** Jennifer Jiang, Mikhail V. Keniya, Anusha Puri, Xueying Zhan, Jeff Cheng, Huan Wang, Gigi Lin, Yun-Kyung Lee, Nora Jaber, Yasmine Hassoun, Erika Shor, Zheng Shi, Sang-Hyuk Lee, Min Xu, David S. Perlin, Wei Dai

## Abstract

Fungal plasma membrane proteins represent key therapeutic targets for antifungal agents, yet their structure and spatial distribution in the native context remain poorly characterized. Herein, we employ an integrative multimodal approach to elucidate the structural and functional organization of plasma membrane protein complexes in *Candida glabrata*, focusing on prominent and essential membrane proteins, the polysaccharide synthase β-(1,3)-glucan synthase (GS) and the proton pump Pma1. Cryo-electron tomography (cryo-ET) and live cell imaging reveal that GS and Pma1 are heterogeneously distributed into distinct plasma membrane microdomains. Treatment with caspofungin, an echinocandin antifungal that targets GS, alters the plasma membrane and disrupts the native distribution of GS and Pma1. Based on these findings, we propose a model for echinocandin action that considers how drug interactions with the plasma membrane environment lead to inhibition of GS. Our work underscores the importance of interrogating the structural and dynamic characteristics of fungal plasma membrane proteins *in situ* to understand function and facilitate precisely targeted development of novel antifungal therapies.

## Main

Invasive fungal infections affect more than a billion people and account for about 1.7 million deaths worldwide per year^1^. The most common opportunistic human fungal pathogens, particularly among individuals with pre-existing conditions, include *Candida* spp., *Aspergillus* spp. and *Cryptococcus* spp.^2,3^. Despite posing significant threats to public health, fungal infections remain neglected and poorly addressed. The most common FDA-approved antifungal agents in clinical use either target the fungal-specific ergosterol in the plasma membrane or its biosynthesis (polyenes and azoles, respectively), or disrupt cell wall integrity (echinocandins)^4,5^. However, the effectiveness of these antifungals has been increasingly compromised by drug resistance^6,7^. Therefore, improvement in existing antifungal compounds and identification of novel therapeutic strategies are urgently needed to combat resistant fungal pathogens.

The fungal plasma membrane and associated proteins have increasingly been recognized as promising targets for antifungal therapy. Recent advances in structural determination using single particle cryo-electron microscopy (cryo-EM) have provided structural insights into several fungal plasma membrane proteins, the H^+^-ATPase Pma1, chitin synthase, and β-(1,3)-glucan synthase (GS) ^8–11^. GS is responsible for synthesis of glucans within the fungal cell wall and is the target for the FDA-approved echinocandin drug class^12,13^. Although the high-resolution structure of GS has revealed the structural basis of glucan synthesis (PDB ID 7XE4^11^), the molecular mechanisms underlying echinocandin inhibition remain elusive. While clinical studies have linked acquired echinocandin resistance to amino acid substitutions in highly conserved hotspot regions of the catalytic subunit of GS, encoded by *FKS1* and *FKS2* genes^14^, recent studies have shown that modifications of the lipid compositions can affect GS susceptibility to echinocandins, indicating that the native membrane environment plays a role in GS structural integrity and its interactions with echinocandins^11,15,16^. Investigating the structure and dynamics of fungal plasma membrane proteins within their native membrane environment is crucial to understanding their roles in fungal physiology and their potential as antifungal drug targets.

In this work, we applied a multidisciplinary approach encompassing cryo-electron tomography (cryo-ET) with quantitative proteomics, biochemical, and biophysical analyses to curate a structural atlas of the fungal plasma membrane proteome *in situ.* We demonstrate that two fungal plasma membrane proteins (*i.e.* GS and Pma1) are organized in functional microdomains within the plasma membrane. Echinocandin treatment modifies the properties of the plasma membrane, which correlates with perturbed spatial distribution of GS and Pma1. Our study characterizes the functional distribution of prominent fungal plasma membrane proteins and provides critical insights into their implications in echinocandin action.

## Results

### An integrative workflow towards visual proteomics of the fungal plasma membrane

To enable *in situ* structural studies of fungal plasma membranes, we applied an integrative framework that combines proteomics, microbiological, and biophysical analyses with multimodal bioimaging techniques (**Fig. 1**). We first isolated membrane ghosts from spheroplasts of *C. glabrata* cells to perform proteomic analysis to define the protein composition of the fungal plasma membrane. Combining quantitative mass spectrometry with cryo-electron tomography (cryo-ET) and deep-learning based annotation enabled molecular characterization of the fungal plasma membrane landscape *in situ*. We performed antifungal susceptibility testing to determine the minimum inhibitory concentration (MIC) of caspofungin (CSF), an echinocandin drug. Micropipette aspiration on intact spheroplasts assessed whether CSF modifies the biophysical properties of the plasma membrane. Integration of these quantitative and qualitative studies provides a robust, comprehensive framework for unraveling the functional distribution of fungal plasma membrane proteins and the integral role of the membrane environment in echinocandin drug action.

**Figure 1.**
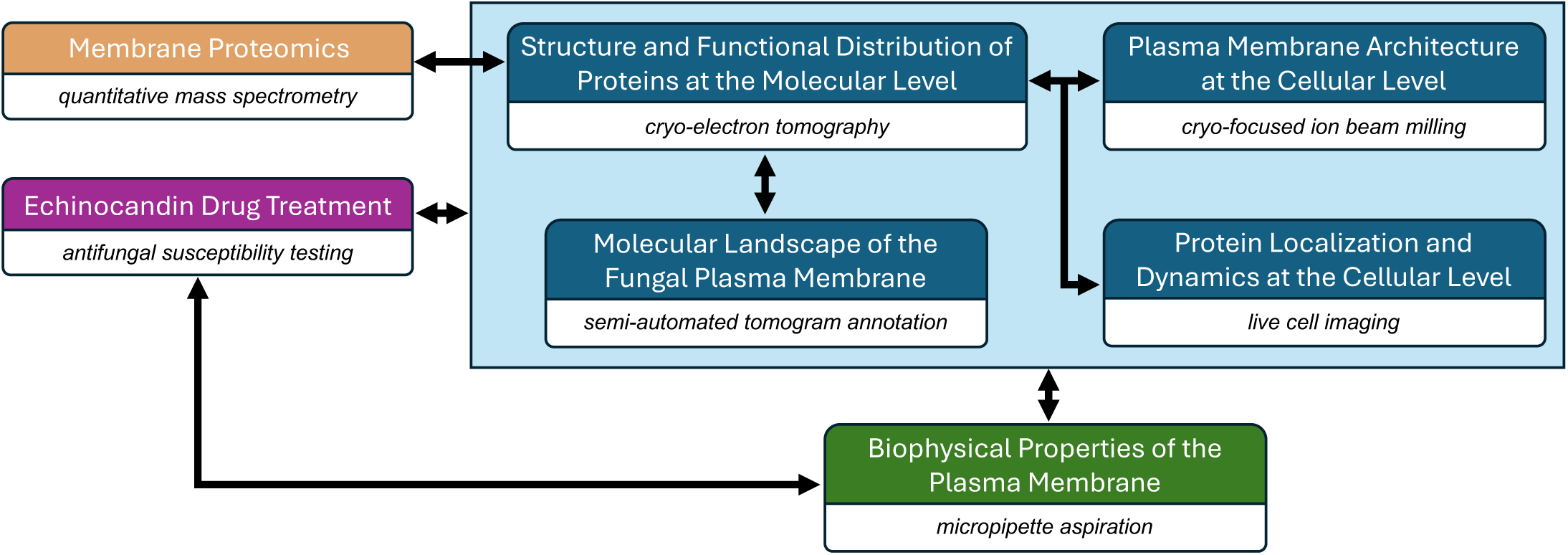
A multidisciplinary approach that enables visual proteomics of the fungal plasma membrane. Integration of multimodal bioimaging techniques (blue boxes) with quantitative proteomics (orange) microbiological (magenta) and biophysical (green) analyses provides a rich toolkit to characterize the functional distribution pattern of fungal plasma membrane proteins in their native environment and dissect echinocandin mechanism of action. Black connecting arrows indicate correlation of results generated from different experimental methods.

To prepare plasma membranes for proteomics and structural studies, we subjected *C. glabrata* cells to enzymatic digestion to remove the cell wall. These spheroplasts were gently ruptured through hypotonic shock to generate crude membranes. Mass spectrometry analysis of these crude membranes identified 3,905 proteins, encompassing 74% of the predicted *C. glabrata* proteome. These proteins are associated with the nucleus, mitochondrial function, ribosome biogenesis and other organelles. Extensive coverage of the proteome indicates that our sample preparation method effectively preserves membranes and associated membrane protein complexes in a close-to-native state for subsequent *in situ* structural investigation.

Overall, the most abundant plasma membrane proteins identified were involved in eisosome formation, cell wall biosynthesis, and cellular transport (**Supplementary Table 1**). To estimate the number of functional complexes expected to be observed on the plasma membrane, we normalized the relative abundance of these proteins to known oligomeric state (**Fig. 2A**). We focused on integral membrane proteins or multi-protein complexes with aggregate molecular weights larger than 100 kDa, since those below this size threshold may not be detectable by cryo-ET. Following this criterion, Fks1, the catalytic subunit of GS, is the most abundant membrane protein that can be detected within the plasma membrane. A single particle cryo-EM structure of Fks1 from *Saccharomyces cerevisiae* suggests that GS is present as a monomer and has a significant cytoplasmic domain (PDB ID 7XE4^11^). The Aqy1 homotetramer, a transmembrane water channel, ranked as the second most abundant membrane protein, followed by Inp53 and Osh2, which are involved in endocytosis and lipid transport, respectively. The fungal H^+^-ATPase Pma1 is another prevalent plasma membrane protein and is shown to assemble into a functional hexameric complex^8,9^. Detection of membrane-embedded proteins by cryo-ET depends on the presence of extramembrane domains. Given that Aqy1, Inp53 and Osh2 are largely embedded within the plasma membrane and lack significant extra membranous projections, these proteins may not be readily detectable by cryo-ET^17,18^. In contrast, Fks1 and Pma1 are sufficiently large or exhibit distinct morphology, which make it possible to identify them in cellular tomograms. Consistent with our proteomics analysis, we observed diverse membrane structures from tomograms, including plasma membranes and those derived from mitochondria and endoplasmic reticulum (ER). We differentiated plasma membranes from other organelle membranes based on overall membrane morphological characteristics and distinctive structural features. For example, rough endoplasmic reticulum (RER) membranes are associated with ribosomes, while mitochondrial membranes are characterized by the presence of distinct cristae, ATP synthases and pyruvate dehydrogenase complexes (**Supplementary Figure 1**).

**Figure 2.**
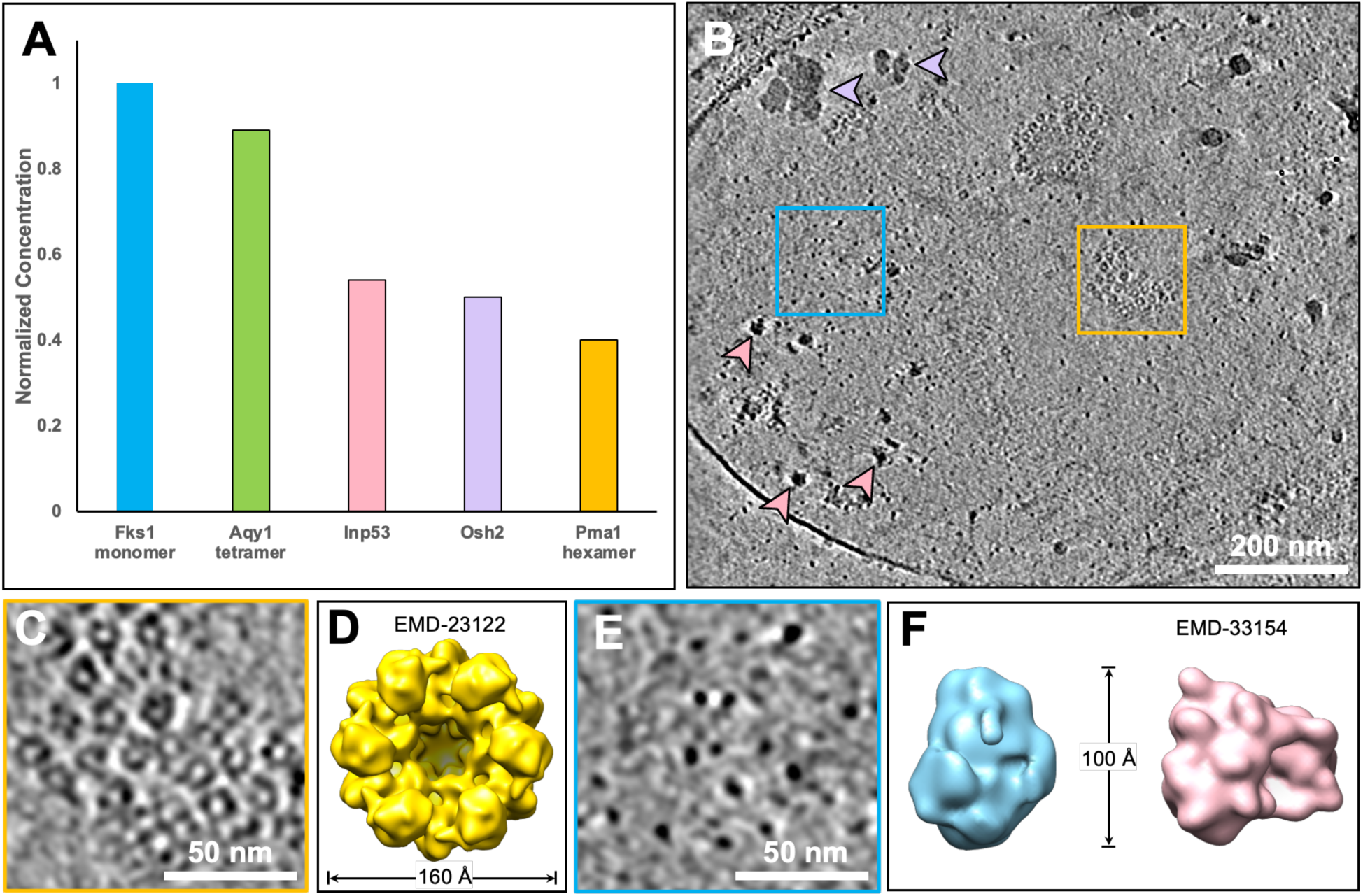
Proteomics analysis and cryo-ET studies characterize the abundance and spatial distribution of fungal plasma membrane proteins in *Candida glabrata*. (**A**) The relative abundance levels of plasma membrane proteins from proteomic analysis of *C. glabrata* membrane ghost preparations with molecular weights larger than 100 kDa, normalized to their known oligomeric state. The protein abundance is expressed relative to the level of Fks1. (**B**) Slice view of a fungal plasma membrane tomogram containing diverse membrane and cytosolic structure features. Purple arrowheads indicate glycogen granules and pink arrowheads depict ribosomes. Scale bar, 200 nm. (**C**) Zoomed-in slice view showing rosette-like particles corresponding to Pma1 hexamers boxed in (B). Scale bar, 50 nm. (**D**) Subtomogram average of Pma1 hexamer (EMD-23122). (**E**) Zoomed-in slice view showing particles corresponding to GS densities boxed in (B). Scale bar, 50 nm. (**F**) Top view of the GS subtomogram average (blue) and the cryo-EM structure of the Fks1 monomer from *Saccharomyces cerevisiae* (pink: EMD-33154)^11^ filtered to 20 Å resolution.

Tomograms collected on fungal plasma membranes revealed a diversity of macromolecular complexes (**Fig. 2B**). The most notable structural features are rosette-like densities with a diameter of approximately 160 Å (**Fig. 2C-D; Supplementary Movie S1**). These structures are often present in clusters of varying sizes and correspond to the overall size and morphology of Pma1 oligomers^8,9^. Additionally, Pma1 clusters appear to induce noticeable membrane depressions, which are clearly visible where side views of Pma1 complexes appear as V-shaped protrusions on the plasma membrane (**Supplementary Fig. 2B, yellow arrows**).

Within the plasma membrane tomograms, we also observed oval-shaped densities (**Fig. 2B, blue box; Supplementary Movie S1**). Given the prevalence of GS in our proteomics analysis, we hypothesize that these densities correspond to GS monomers. The high-resolution cryo-EM structure of GS monomer exhibits a bulky cytoplasmic domain that measures approximately 100 Å in width and 45 Å in height^11^. Subtomogram averaging of these elongated densities confirm that they exhibit overall shape and dimensions consistent with the structure of the cytosolic region of the GS monomer (**Fig. 2E, F; Supplementary Fig. 3**). Due to the preferred orientation of plasma membranes on the EM grid, top views of GS were more predominant, resulting in a poorly resolved transmembrane region.

Other cellular features were also clearly visible in our cellular tomograms. Large, globular-shaped densities distributed throughout the tomograms correspond to ribosomes (**Fig. 2B, pink arrowheads**; **Supplementary Fig. 2**). We also observed twisted, actin filaments (**Supplementary Fig. 2A-B, blue arrowheads**), which have been reported to associate with GS complexes and play a crucial role in maintaining cell wall organization and remodeling^19^. Clover-like granules with diameters of 50–70 nm represent aggregates of glycogen granules that have been previously reported in fungi (**Fig. 2B**; **Supplementary Fig. 2C, purple arrowheads**)^20–22^. These observations from our tomography study, in conjunction with the proteomics analysis, confirm that our sample was preserved in a near-native state.

### Fungal plasma membrane proteins segregate into distinct microdomains in the plasma membrane

To elucidate the spatial interrelationships of the membrane complexes, we employed convolutional neural network (CNN)-based annotation to identify subcellular features within fungal plasma membrane tomograms^23^. Ribosomes and Pma1 hexamers were selected as targets due to their distinct morphology and substantial size, allowing us to use manual annotation as a benchmark to calculate evaluation metrics. The CNN-based approach demonstrated robust performance in detecting ribosomes, achieving an F1 score of 0.796 (**Supplementary Table 2**). The relatively high F1 score indicates that the automated annotation of ribosomes achieved a balance between accuracy and sensitivity. Annotation of ribosomes was independent of isosurface as the performance remained consistent across several isosurface threshold values. In contrast, annotation of Pma1 hexamers was less effective with an F1 score of 0.498 and was significantly influenced by the isosurface threshold. Lowering the threshold improved the annotation performance of Pma1 relative to higher isosurface thresholds. Compared to ribosomes, the lower performance of Pma1 annotation and its sensitivity to isososurface settings may be attributed to the inherent challenges of detecting smaller membrane-embedded proteins. We opted for a lower isosurface setting to maximize the detection of membrane protein targets.

Annotation and mapping of Pma1 and GS complexes within the plasma membrane allowed further analysis of their spatial distribution. To ensure accuracy, we manually inspected the results of the automated annotation to eliminate obvious false positives and to include missed structures. We calculated the density of annotated Pma1 clusters and GS complexes using plasma membrane tomograms with high contrast. On average, Pma1 clusters occupy 17% of the fungal plasma membranes with a density of 264 hexamers per square micrometer of plasma membrane area. GS particles are more densely distributed, with 1,415 monomers per square micrometer. The ratio of GS monomers to Pma1 hexamers in the plasma membrane tomograms is 5.4. In *C. glabrata*, the catalytic subunit of GS is encoded by two homologous genes, *FKS1* and *FKS2*^24^. Given this information, our annotation aligns well with the proteomics analysis, which revealed a ratio of 3.3 Fks1/2 proteins to Pma1.

Pma1 hexamers arrange into dense, crystalline-like microdomains within the plasma membrane (**Fig. 3A, Supplementary Movie 1**). Recent studies show that fungal plasma membranes organize into two main membrane microdomains: the MCP (membrane compartment occupied by ATPase Pma1) and the MCC (membrane compartment occupied by arginine permease Can1). MCCs are static membrane structures that accumulate various integral membrane proteins involved in nutrient transport, response to stress conditions and regulation of lipid biosynthesis^25–28^. Enriched in Pma1, MCPs form large, network-like structures within the plasma membrane and play a role in dynamic vesicular trafficking^29,30^. The heterogenous distribution of Pma1 complexes in our annotated tomogram confirm the presence of membrane microdomains in fungal plasma membranes.

**Figure 3.**
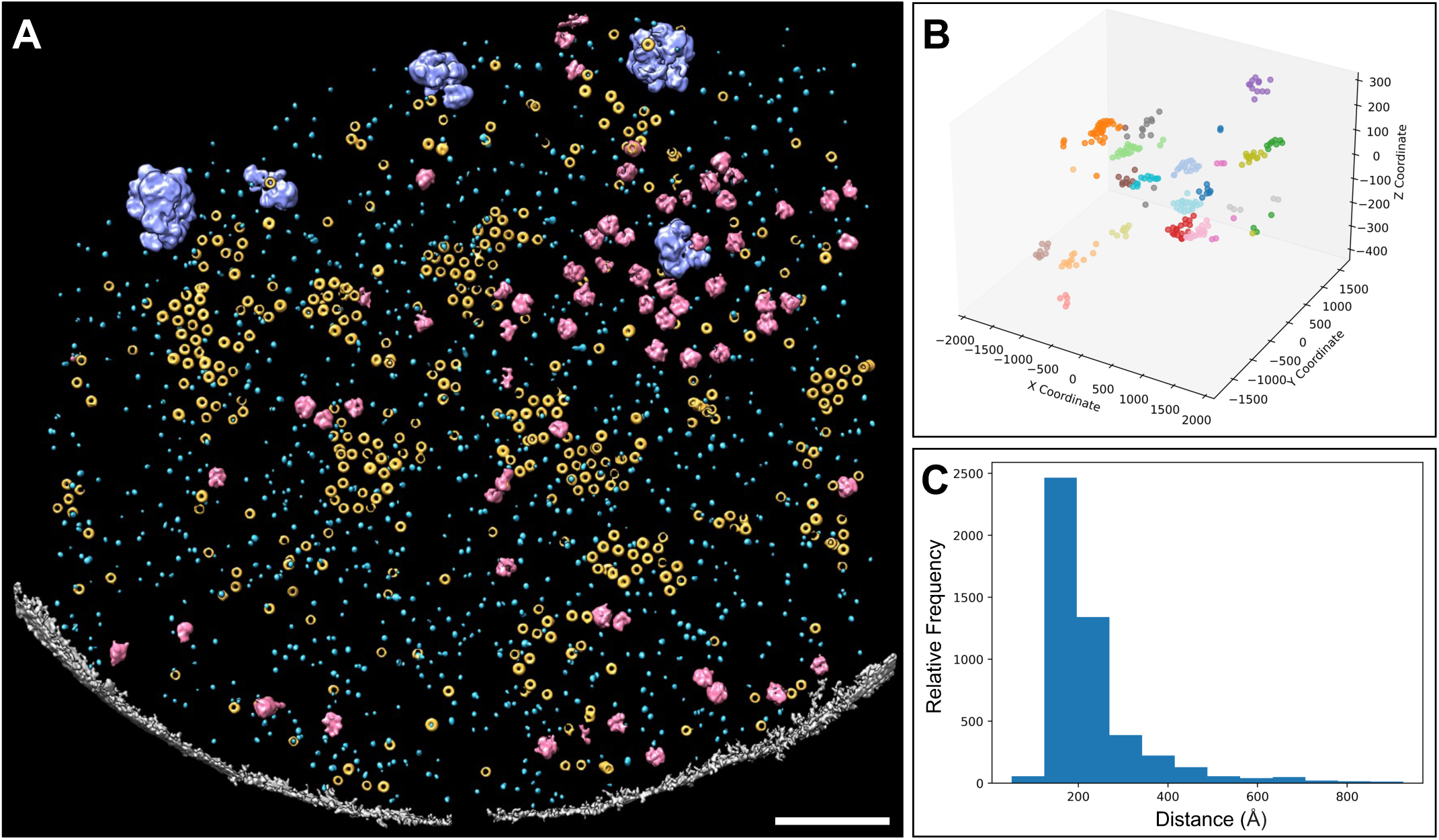
Semi-automated CNN-based annotation revealed the molecular landscape of the fungal plasma membrane. (**A**) Annotation of various structural features in a plasma membrane tomogram. Neural networks were trained independently to recognize Pma1 hexamers (yellow), GS (light blue), ribosomes (pink) and glycogen granules (purple) in context of the plasma membrane (gray). Scale bar, 200 nm. (**B**) 3D scatter plot capturing the spatial distribution of individual Pma1 hexamers, represented as dots, within distinct clusters from a representative plasma membrane tomogram. (**C**) Histogram showing nearest neighbor distance of Pma1 hexamers within clusters in plasma membranes.

To further explore membrane microdomain organization in intact cells, we utilized cryo-focused ion beam milling to thin vitrified *C. glabrata* cells and performed cryo-ET imaging on yeast lamellae. Tomograms of intact cells displayed several furrow-like invaginations on the plasma membrane that have a length in the range 40–70 nm of and varying depth of 160–250 nm (*n* = 13), characteristic of MCC structures (**Supplementary Fig. 4, teal arrowheads; Supplementary Movie S2**)^31^. We also observed densities of membrane protein complexes interspersed along the invaginated membranes. Taken together, cryo-lamellae of intact cells and tomogram annotation support the concept that fungal plasma membranes partition membrane proteins into distinct microdomains.

Given that Pma1 is the landmark protein of MCPs, further characterizing its spatial distribution may provide insights into the functional organization of the plasma membrane. A 3D scatter plot of Pma1 complexes from a representative plasma membrane tomogram reveal variations in the size and shape of distinct clusters (**Fig. 3B**). Nearest neighbor analysis indicated that Pma1 hexamer within the clusters follows a Gaussian distribution with a narrow peak at 180 Å, suggesting compact packing of Pma1 complexes (**Fig. 3C**). We then manually identified the number of Pma1 clusters within our plasma membrane tomograms based two criteria: 1) Pma1 hexamers within a cluster are positioned approximately 160–170 Å apart from one another and 2) a minimum of four particles are required to constitute a cluster. To validate our manual clustering outcome, we applied Gaussian Mixture Models (GMM) and *k*-means clustering on annotated Pma1 complexes to estimate the number of clusters. The number of Pma1 clusters determined by both GMM and *k*-means clustering methods closely aligned with manual assignment of clusters, further confirming that Pma1 arrange into clusters in the plasma membrane (**Supplementary Table 3**).

While GS complexes in annotated tomograms also exhibit some level of higher-order organization, assessing their spatial distribution is challenging due to their overall denser and variable distribution. To alternatively characterize the subcellular distribution of GS *in vivo* with fluorescence microscopy, we engineered a strain that expresses Fks1 tagged with the yellow fluorescent protein (YFP) at the N-terminus (IGCg1 strain). This strain grows normally and has a similar echinocandin susceptibility profile as the wild type strain (CBS138), indicating that addition of the N-terminal YFP tag does not affect GS activity (**Supplementary Figure 5**). Distinct YFP-FKS1 punctae with an average diameter of half a micron [0.51 ± 0.083 µm (1 SD), *n* = 31] were observed along the surface of these cells **(Fig. 4C**), suggesting that GS are heterogeneously distributed and form distinct microdomains within the plasma membrane. Staining these cells with aniline blue dye showed that the glucan content within the cell wall appeared heterogenous, as indicated by the non-uniform fluorescent pattern along the cell periphery (**Fig. 4A-C**). Notably, nascent budding yeasts exhibited stronger aniline blue staining, consistent with the fact that β-(1,3)-glucans synthesis actively occurs at sites of budding (**Fig. 4B, white arrows**).

**Figure 4.**
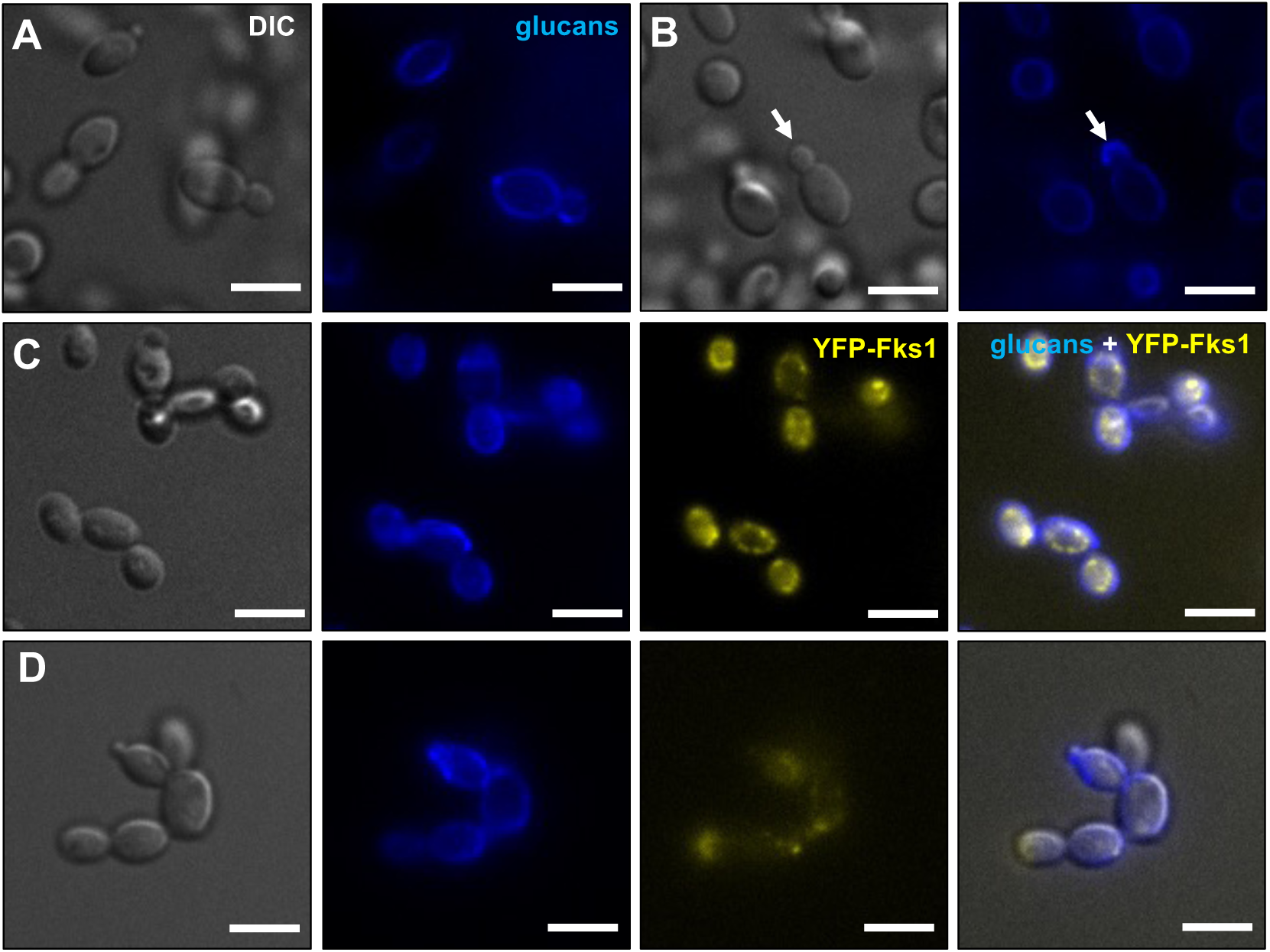
The heterogeneous distribution of glucan synthase (GS) within *Candida glabrata* cells is altered following caspofungin (CSF) treatment. (**A-B**) CBS138 (wild type) and (**C-D**) IGCg1 (YFP-Fks1-expressing) strains stained with glucan-specific aniline blue to reveal the *in vivo* glucan levels within the cell wall. Localization of GS within (**C**) untreated and (**D**) CSF-treated IGCg1 cells. White arrows indicate nascent budding cells. Scale bars, 5 µm.

### Caspofungin treatment alters the spatial localization of GS and Pma1

GS is the target for FDA-approved echinocandin and enfumafungin class of antifungals, which compromise the cell wall integrity and renders cells susceptible to osmotic shock. In clinical settings, resistance to echinocandins is primarily linked to mutations in three hotspot regions of GS encoded by *FKS* genes^14,32,33^. GS membrane topology analysis and the high-resolution cryo-EM structure of GS mapped these hotspot regions to the extracellular side of the transmembrane domain suggesting these hot spot regions form the echinocandin drug binding pocket for echinocandin drugs^11,34^. Interestingly, in the high resolution cryo-EM structure of GS, multiple lipid molecules are shown associated with these hotspot regions, suggesting that these local lipids may be involved in echinocandin interactions with GS^11^. To explore the relationship between echinocandins and the lipid membrane environment, we challenge IGCg1 cells with caspofungin (CSF), the first-in-class echinocandin antifungal. Fewer and less distinct YFP-Fks1 punctae were observed along the cell periphery compared to untreated cells, suggesting that CSF treatment disrupts the organization of GS into microdomains within the plasma membrane (**Fig. 4C-D**).

To investigate whether CSF exposure alters the spatial arrangement of fungal membrane proteins at the molecular level, we conducted cryo-ET on plasma membranes generated from CSF-treated spheroplasts. Our analysis included a strain that overexpresses Fks1 (KH238) to investigate whether increased GS abundance may affect echinocandin action. In the Fks1-overexpressing strain, Pma1 clusters are typically more prevalent and larger, likely due to membrane crowding from overexpression of GS^35^.

CSF treatment altered the clustering dynamics of Pma1 hexamers within plasma membranes from both the wild type and Fks1-overexpressing cells. Specifically, Pma1 complexes become more dispersed within the plasma membrane (**Fig. 5**; **Supplementary 3**). Annotation of a representative tomogram of CSF-treated wild type plasma membrane revealed the disruption of Pma1 clustering (**Supplementary Fig. 6**, **Supplementary Movie 3**). Using *k-*means clustering, we observed that the average intra-cluster distance and cluster radius of Pma1 hexamers in wild type membranes increased by 1.7-fold after CSF treatment (**Fig. 5G, H**; **Supplementary Table 4**). A similar trend was observed with Fks1-overpressing plasma membranes following drug exposure with a 1.9-fold and 2.1-fold increase in the average intra-cluster distance and average cluster radius, respectively. Smaller, tightly-packed Pma1 clusters were also present in CSF-treated Fks1 overexpressing plasma membranes (**Fig. 5E, F**). In addition, CSF-treated plasma membranes often exhibit deformities in the membrane, suggesting that CSF exposure impacts the plasma membrane ultrastructure.

**Figure 5.**
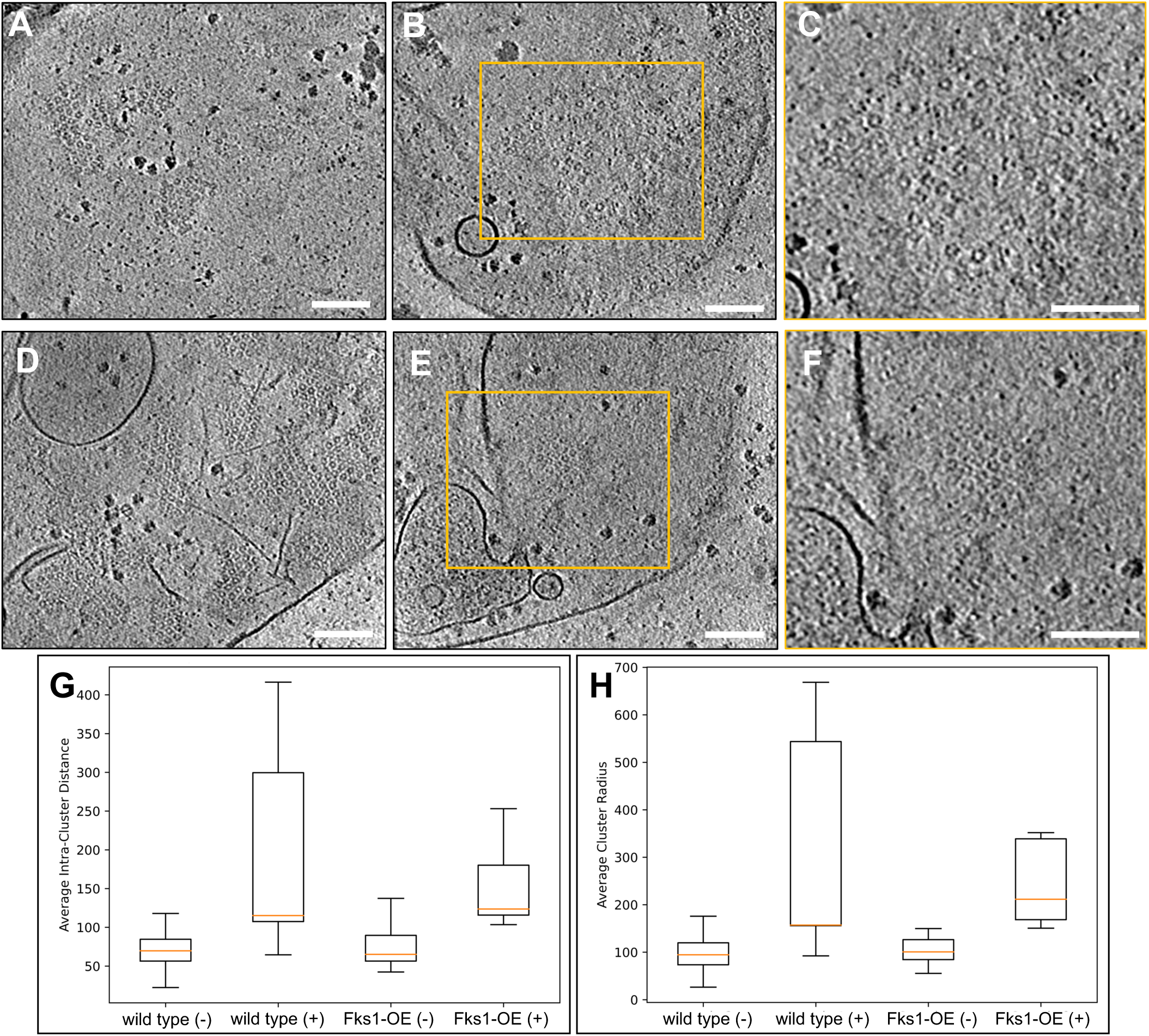
Caspofungin (CSF) perturbed the spatial distribution patterns of Pma1 in *Candida glabrata* plasma membranes. (**A-B**) Slice views of an untreated (**A**) and a CSF-treated (**B**) wild type plasma membrane tomogram. (**C**) Zoomed-in view of dispersed Pma1 hexamers boxed in (**B**). (**D-E**) Slice views of an untreated (**D**) and a CSF-treated (**E**) KH238 plasma membrane tomogram. (**F**) Zoomed-in view of a small Pma1 cluster highlighted in (**E**). (**G**) Intra-cluster distance analysis (nm) of Pma1 clusters in wild type and Fks1-overexpressing plasma membranes. (**H**) Analysis of average cluster radius (nm) in wild type and Fks1-overexpressing plasma membranes before and after CSF treatment. wild type (-): *n* = 14; wild type (+): *n* = 16; Fks1-OE (-): *n* = 19; Fks1-OE (+): *n* = 7. (-/+) indicates without or with CSF treatment. Scale bars, 100 nm.

We also examined the spatial distribution of Pma1 hexamers in plasma membranes from a strain with a mutation in the FEN1 gene encoding a fatty acid elongase (MVKCg9). This strain exhibits defective sphingolipid biosynthesis and a 6-fold decrease in CSF susceptibility (**Supplementary Figure 5**). Notably, even in the absence of CSF treatment, plasma membranes from Δfen1 mutant appear to exhibit rough and textured membrane morphology (**Supplementary Fig. 7A**). Pma1 hexamers within Δfen1 plasma membranes were also predominantly dispersed, a distribution pattern similar to that observed in CSF-treated CBS138 and KH238 plasma membranes (**Supplementary Fig. 7A-C**).

Taken together, our findings suggest that CSF disrupts the native spatial organization of fungal plasma membrane proteins, implying that echinocandin-mediated inhibition of glucan synthesis involves alterations to the overall membrane environment.

### Caspofungin treatment alters the properties of the spheroplast membrane

Echinocandins are characterized by a cyclic hexapeptide core with a lipophilic tail. Given our observations that CSF perturbs plasma membrane ultrastructure and induces spatial reorganization of membrane proteins, it is plausible that echinocandins can directly interact with and modify the properties of the plasma membrane. To explore this hypothesis, we generated spheroplasts from both the wild type and Δfen1 mutant strains to evaluate potential changes in the biophysical properties of the plasma membrane following CSF treatment (**Fig. 1**). Compared to yeast cells with intact cell wall, spheroplasts generally exhibit a spherical morphology with an enlarged diameter of 5–6 µm (**Fig. 6A-B**). Notably, Δfen1 spheroplasts were slightly smaller than wild type spheroplasts, likely due to deficiency in lipid metabolism affecting plasma membrane synthesis or recycling^15,36^. However, treatment with CSF did not affect the spheroplast size in either strain (**Fig. 6C**).

**Figure 6.**
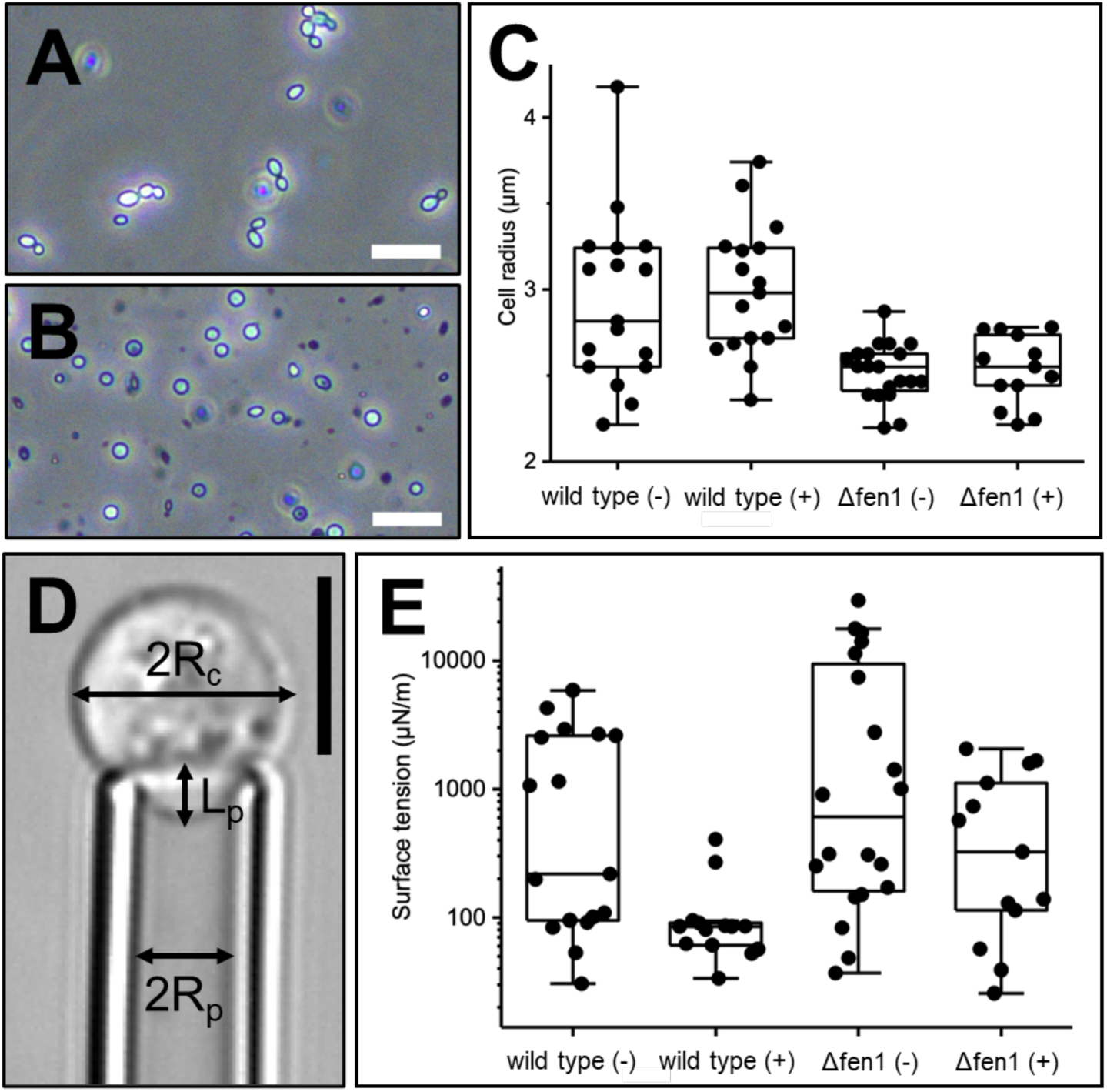
Caspofungin (CSF) treatment altered the biomaterial properties of the plasma membrane. (**A-B**) Light microscopy images of *Candida glabrata* cell-walled cells (**A**) and cell wall less-spheroplasts (**B**). Scale bars, 20 µm. (**C**) Distribution of wild type and Δfen1 mutant spheroplast size in the absence and the presence of CSF treatment. (**D**) Micropipette aspiration was used to measure the surface tension of spheroplast. R_c_: cell radius; R_p_: micropipette radius; L_p_: length of the portion aspirated into the micropipette. Scale bar, 5 µm. (**E**) Surface tension of wild type and Δfen1 spheroplasts in 1M sorbitol, with and without CSF treatment. wild type (-): *n* = 17; wild type (+): n = 14; Δfen1 (-): *n* = 20; Δfen1 (+) : *n* = 13. (-/+) indicates without or with CSF treatment.

We then performed micropipette aspiration (MPA) to evaluate the effect of CSF on spheroplasts from wild type and Δfen1 mutant strains. We determined the surface tension by measuring the degree of membrane deformation in response to stepwise increase in aspiration pressure (**Fig. 6D-E; Supplementary Movie 4 and 5**). The median surface tension of wild type and Δfen1 spheroplasts decreased by 2.6- and 1.9-fold, respectively, following CSF treatment (**Fig. 6F**). In conjunction with tomography data, our biophysical analyses support the notion that echinocandins directly interact with and modify the local lipid microenvironment surrounding fungal plasma membrane protein complexes, and thereby alter their functional spatial distribution.

## Discussion

The fungal plasma membrane and its resident proteins have been intensively studied as potential targets for antifungal therapy. Recent advancements in cryo-EM single particle analysis have laid the groundwork for structural determination of several key fungal plasma membrane proteins, including Pma1^8,9^, chitin synthase^10^, and GS^11^. These high-resolution structures have provided insights into the molecular mechanisms underlying their role in fungal physiology and underscore the influence of the native membrane environment in regulating protein function and drug interactions. By using an integrative multimodal approach with cryo-ET as the cornerstone technique, we visualize the molecular landscape of the fungal plasma membrane and examine drug-membrane interactions within the cellular context. Our findings demonstrate that prominent fungal plasma membrane proteins, such as Pma1 and GS, are distributed into distinct membrane microdomains. This higher-order organization of membrane proteins into microdomains within the fungal plasma membrane likely contributes to the cell’s ability to perform tightly regulated biological functions in the crowded membrane environment and serves an integral role in fungal infectivity and pathogenicity^37,38^.

Despite decades of clinical and basic research, the molecular mechanisms underpinning echinocandin drug action remain poorly understood. Topological analysis and mutagenesis studies have mapped the hotspot regions implicated in echinocandin resistance onto the extracellular domain of GS^34^, supporting the “direct binding” model for echinocandin action. This model proposes that echinocandins bind to these hotspot regions, induce conformational changes that extend to the active site, and inhibit glucan biosynthesis (**Fig. 7**).

**Figure 7.**
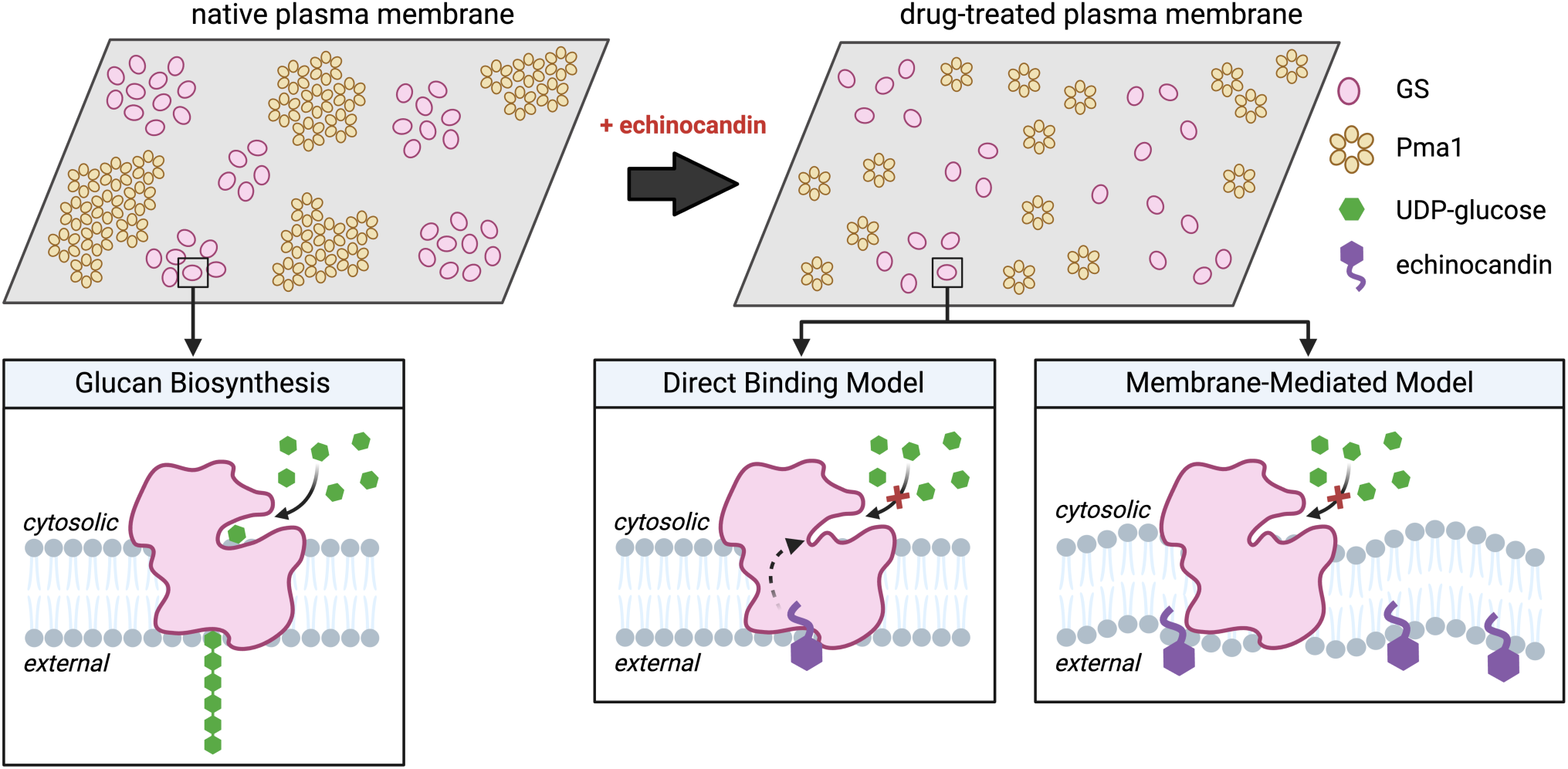
Models for echinocandin drug action. Direct binding model: Echinocandin directly binds to GS and induces conformational changes to the active site of GS, resulting in inhibited glucan synthesis. Membrane-mediated interaction model: Incorporation of echinocandins into the plasma membrane alters the local lipid environment of GS, leading to perturbed GS structure, spatial distribution and functional activity. Created with BioRender.com.

Due to the amphipathic characteristic of echinocandins, it has also been speculated that echinocandins may integrate into the fungal plasma membrane and indirectly inhibit GS function by modifying the local lipid environment. Biochemical studies have indicated that changes in plasma membrane lipid composition, particularly sphingolipid content, affect GS susceptibility to echinocandin inhibition^15,16,36^. Supporting this notion, the Δfen1 mutant, which is defective in sphingolipid biosynthesis, exhibited changes in membrane morphology, membrane protein distribution, and altered sensitivity to CSF treatment. We further demonstrated that exposure to CSF alters the surface tension of the plasma membrane, which is correlated with the perturbed microdomain organization of GS and Pma1. These findings support the theory that echinocandins interact with and modify the membrane environment, and thereby influencing the functional dynamics of embedded membrane proteins (**Fig. 7**). In the cryo-EM structure of GS, ordered lipid density features were identified within the hotspot regions conferring echinocandin-resistance, implicating the critical role of endogenous lipids in the structural and functional dynamics of GS^11^. It is also possible that echinocandins directly interact with lipid molecules within the binding pocket, and thereby modify GS enzymatic activity. Recent docking simulations have suggested that the lipophilic tail of echinocandins may form a ternary complex with these bound lipids^39^. Further studies will be necessary to determine the molecular details of these interactions and to understand how specific lipid molecules, such as sphingolipids and ergosterols, influence echinocandin activity and the development of antifungal resistance.

In this study, we implemented a robust integrative multimodal workflow that combines quantitative proteomics, biochemical and biophysical analyses with cutting-edge bioimaging methods. Our work characterizes the functional distribution pattern of fungal plasma membrane proteins in their native context and provides novel insights into the role of the membrane environment in echinocandin action, opening exciting avenues for the development of more efficacious antifungal strategies.

## Funding and Acknowledgements

We thank Jason T. Kaelber and Emre Firlar at the Rutgers CryoEM & Nanoimaging Facility (RCNF) for their technical assistance in data collection. A portion of this research was supported by NIH grant U24 GM139168 and performed at the Midwest Center for Cryo-ET (MCCET) and the Cryo-EM Research Center in the Department of Biochemistry at the University of Wisconsin. We would like to thank Muyuan Chen for his technical support with EMAN2. We would like to acknowledge Haiyan Zheng at the Biological Mass Spectrometry Facility at Robert Wood Johnson Medical School for performing the mass spectrometry analysis. J. J. and Y. L. were partially supported by the NSF CAREER MCB-2046180. J.J. was also partially supported by a Rutgers University Presidential Graduate Fellowship awarded by the Rutgers Institute for Quantitative Biomedicine. Y.H., E.S., and D.S.P. were partially supported by NIH R01AI109025. X.Z. and M.X. were partially supported by NIH R01GM134020 and NSF CAREER DBI-2238093. S.L. was supported by DOE DE-SC0019313, and H.W. and Z.S. were supported by R35GM147027 and R21DA056322.

## Data Availability

Tomograms of untreated and CSF-treated wild type plasma membranes have been deposited in the EMDataBank under accession codes EMD-45105 and EMD-45106, respectively.

## Supplementary Materials

**Supplementary Table 1.**
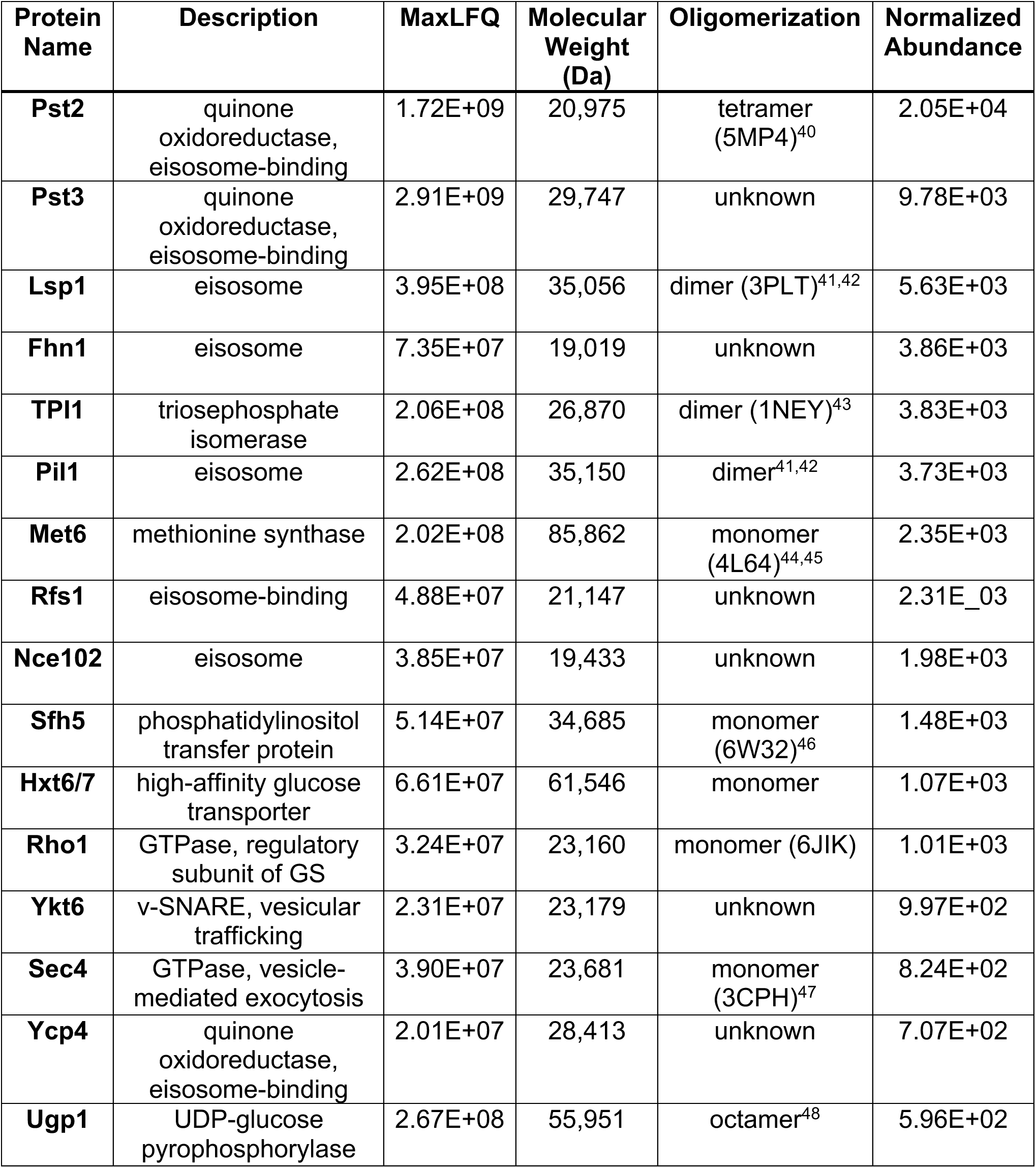
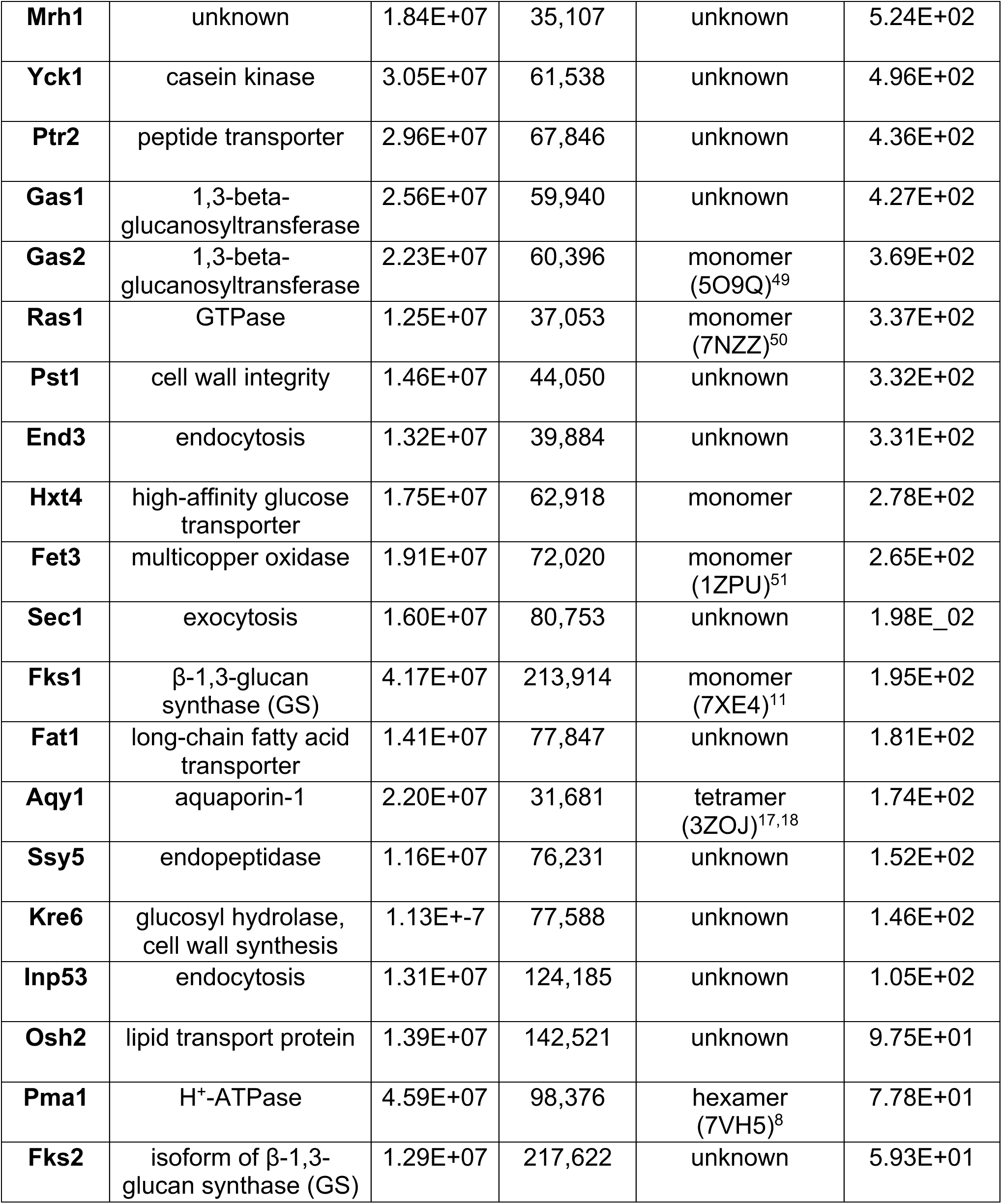
The most abundant plasma membrane proteins identified from *Candida glabrata* crude membranes.

**Supplementary Table 2.** Quantitative evaluation of convolutional neural network (CNN)-based annotation using manual annotation as a benchmark.

|  | manual<br>annotation | EMAN2 annotation |  | true<br>positive | false<br>negative | false<br>positive | F1 score |
| --- | --- | --- | --- | --- | --- | --- | --- |
|  |  | threshold | total picked |  |  |  |  |
| ribosome | 177 | 0.45 | 265 | 172 | 5 | 88 | 0.788 |
|  |  | 0.50 | 263 | 171 | 6 | 86 | 0.788 |
|  |  | 0.55 | 260 | 171 | 6 | 83 | 0.793 |
| Pma1 | 451 | 0.45 | 669 | 222 | 229 | 218 | 0.498 |
|  |  | 0.50 | 607 | 194 | 257 | 156 | 0.484 |
|  |  | 0.55 | 560 | 159 | 292 | 109 | 0.443 |

**Supplementary Table 3.** Quantification of estimated Pma1 clusters in plasma membrane tomograms collected from different strains with and without caspofungin (CSF) treatment as determined by manual labeling, Gaussian Mixture Model (GMM) and *k*-means clustering methods.

| sample<br>(-/+ CSF) | number of<br>tomograms | number of<br>particles | manual<br>estimation | GMM<br>clustering | *k*-means<br>clustering |
| --- | --- | --- | --- | --- | --- |
| CBS138 (-) | 14 | 3289 | 155 | 153 | 155 |
| CBS138 (+) | 16 | 1469 | 52 | 50 | 52 |
| KH238 (-) | 19 | 3289 | 155 | 153 | 155 |
| KH238 (+) | 7 | 455 | 24 | 24 | 24 |

**Supplementary Table 4.** Intergroup comparison showing significance of differences in clustering metrics among the four sample conditions. The Kruskal-Wallis H test yielded *p* < 0.005 for both clustering metrics. Post-hoc multiple comparisons were performed using Dunn’s test. S1 = CBS138 without CSF; S2 = CBS138 with CSF; S3 = KH238 without CSF; S4 = KH238 with CSF.

| clustering metric | Kruskal-Wallis test |  | estimated power | Post-hoc (Dunn's) *p*-value |  |  |  |  |  |
| --- | --- | --- | --- | --- | --- | --- | --- | --- | --- |
|  | H test value | *p*-value |  | S1-S2 | S1-S3 | S1-S4 | S2-S3 | S2-S4 | S3-S4 |
| average cluster radius | 19.803 | **0.0002** | 0.8967 | **0.0036** | 0.586 | **0.0027** | **0.0011** | 0.8561 | **0.0008** |
| average intra-cluster distance | 18.3275 | **0.0004** | 0.9266 | 0.0051 | 0.717 | **0.0023** | **0.0029** | 0.9903 | **0.0014** |

**Supplementary Figure 1.**
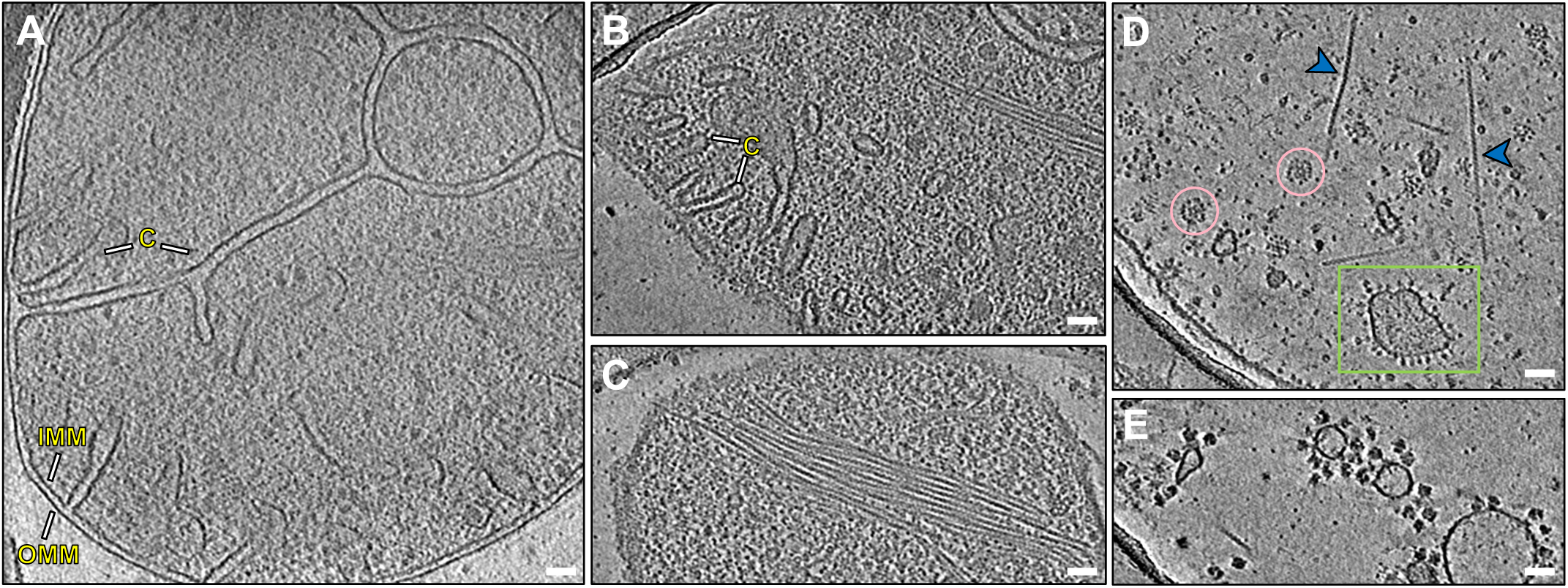
Visualization of diverse cellular organelle features in *C. glabrata*. Slice views of reconstructed cellular tomograms depicting various subcellular features: (**A**) intact mitochondrion with distinct outer membrane (OMM), inner membrane (IMM) and cristae (C), (**B**) mitochondrial membrane with distinct inner membrane cristae (C), (**C**) bundle of actin filaments, (**D**) actin filaments (blue arrowheads), pyruvate dehydrogenase complexes (pink circles) and ATP synthases decorating mitochondrial membrane curvature (green box), and (**E**) membrane-associated ribosomes. Scale bars, 50 nm.

**Supplementary Figure 2.**
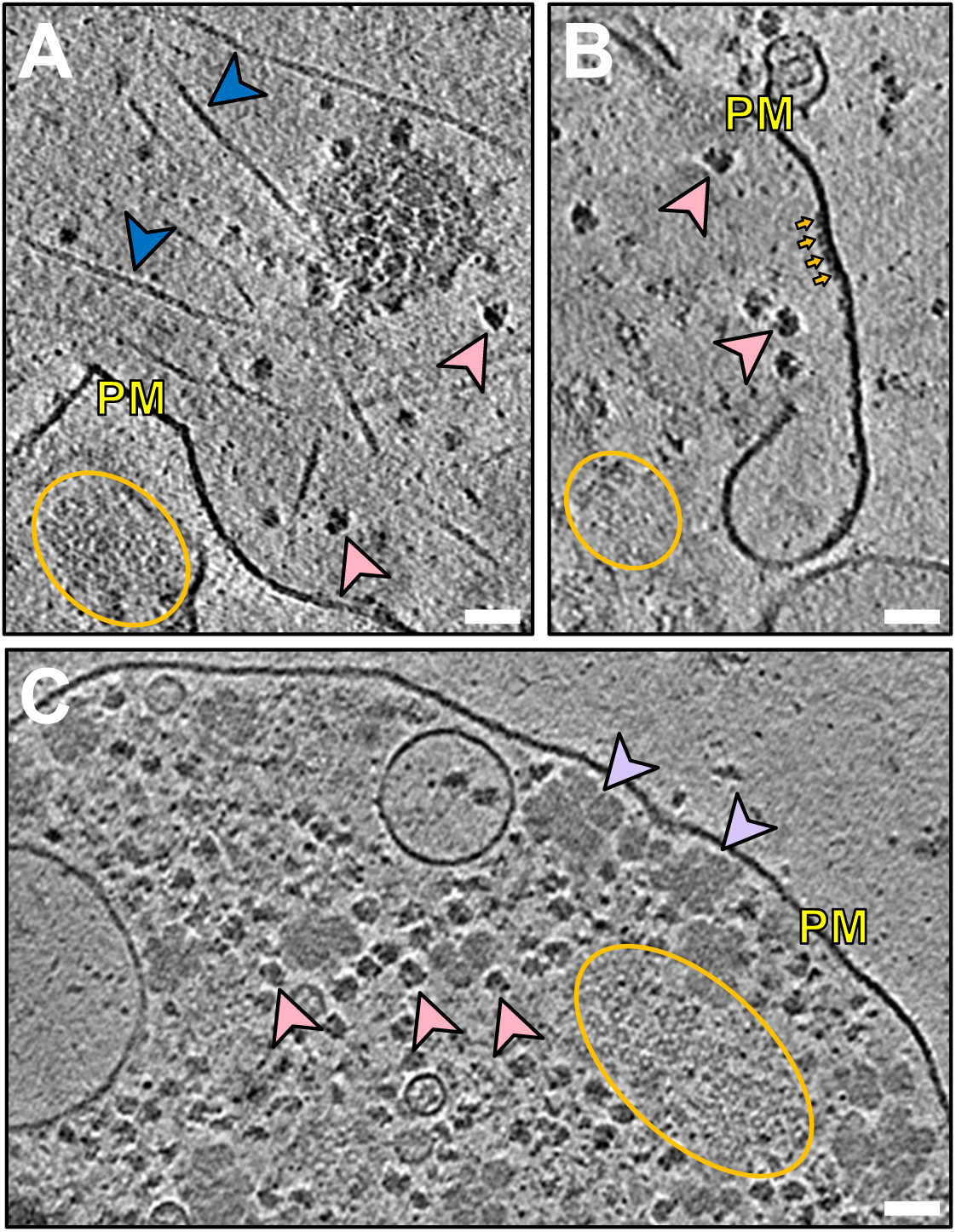
Cryo-ET reveals cellular structures and macromolecular complexes in the fungal plasma membrane. **(A-C)** Slice views of plasma membrane tomograms depicting various structural features including the plasma membrane (PM), cytosolic ribosomes (pink arrowheads), actin filaments (blue arrowheads) (**A**), cluster of Pma1 hexamers (yellow circle), side views of Pma1 hexamers (yellow arrows) (**B**), and glycogen storage granules (purple arrowheads) (**C**). Scale bars, 50 nm.

**Supplemental Figure 3.**
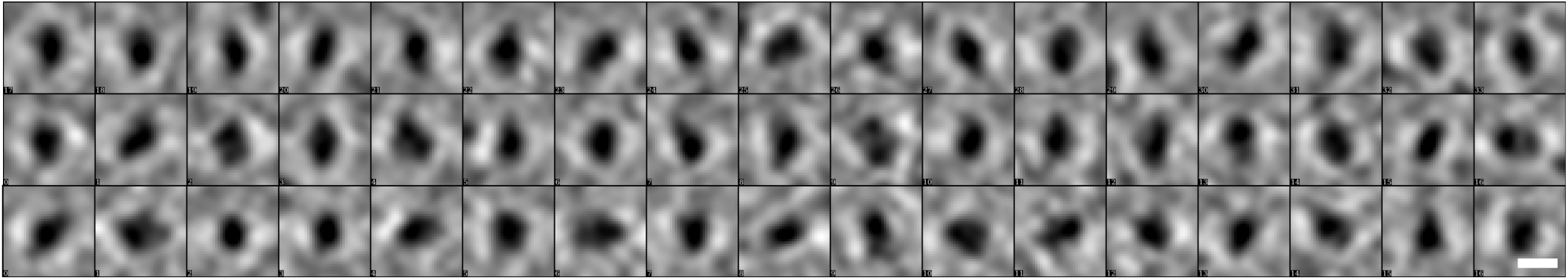
Representative subtomograms of GS. 2D boxing of particles selected from slices of plasma membrane tomograms. Scale bar, 10 nm.

**Supplementary Figure 4.**
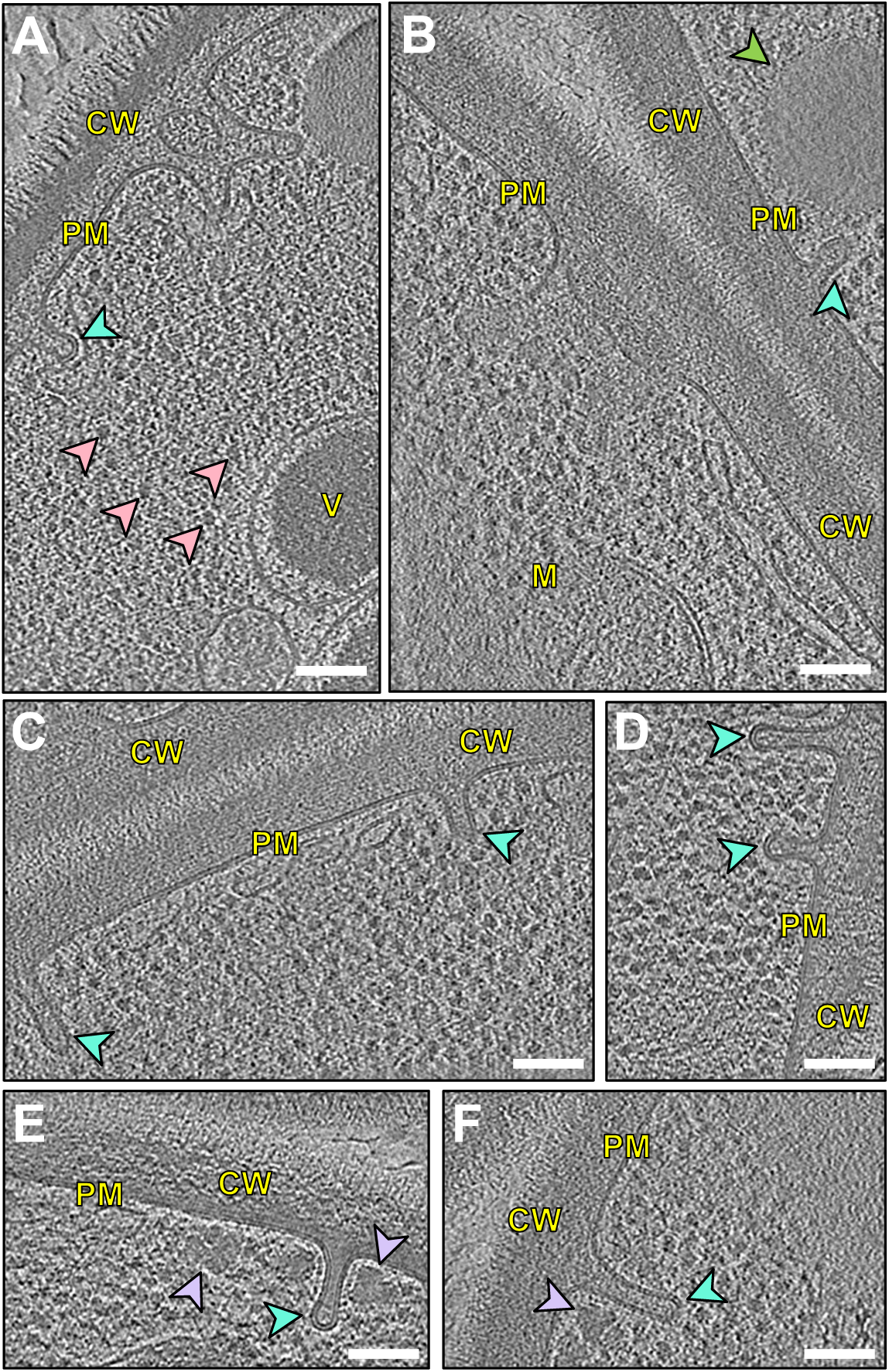
Cryo-ET of lamellae of *Candida glabrata* cells reveals fungal plasma membrane microcompartments. Cellular structures are clearly visible: cell wall (CW), plasma membrane (PM), eisosome (teal arrowhead), ribosome (pink arrowheads), vacuole (V), lipid droplet (green arrowhead), mitochondrion (M), glycogen granule (purple arrowheads). Scale bars, 100 nm.

**Supplementary Figure 5.**
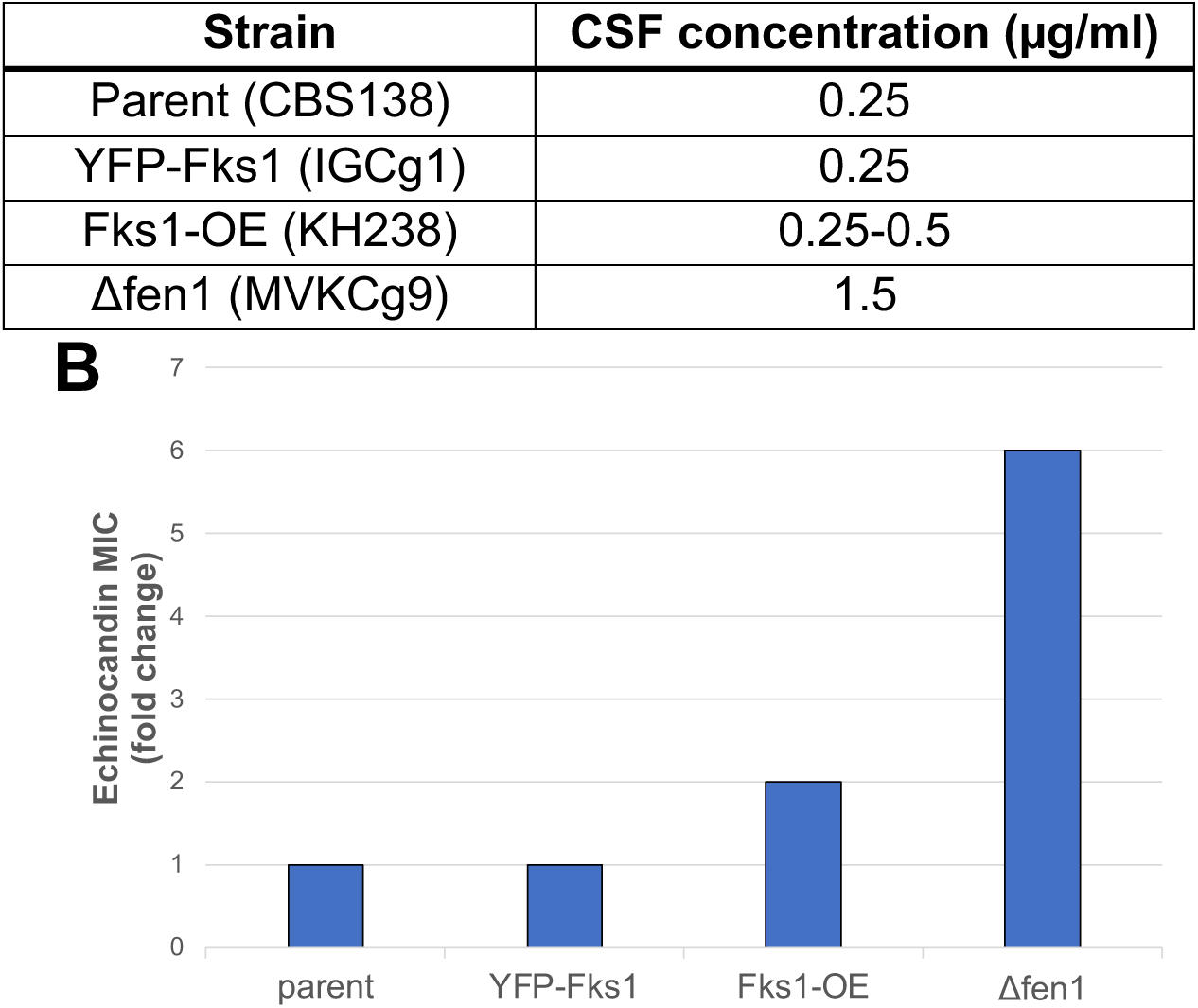
Antifungal susceptibility testing of various *C. glabrata* strains. (**A**) Caspofungin (CSF) minimum inhibitory concentration (MIC) ranges against various *C. glabrata* strains used in this study. The IGCg1 strain expresses YFP-tagged Fks1, and the KH238 strain overexpresses Fks1. MVKCg9 strain harbors a deletion of *FEN1* which encodes a fatty acid elongase. (**B**) CSF susceptibility results expressed as fold changes in the MIC value relative to the parent strain (CBS138).

**Supplementary Figure 6.**
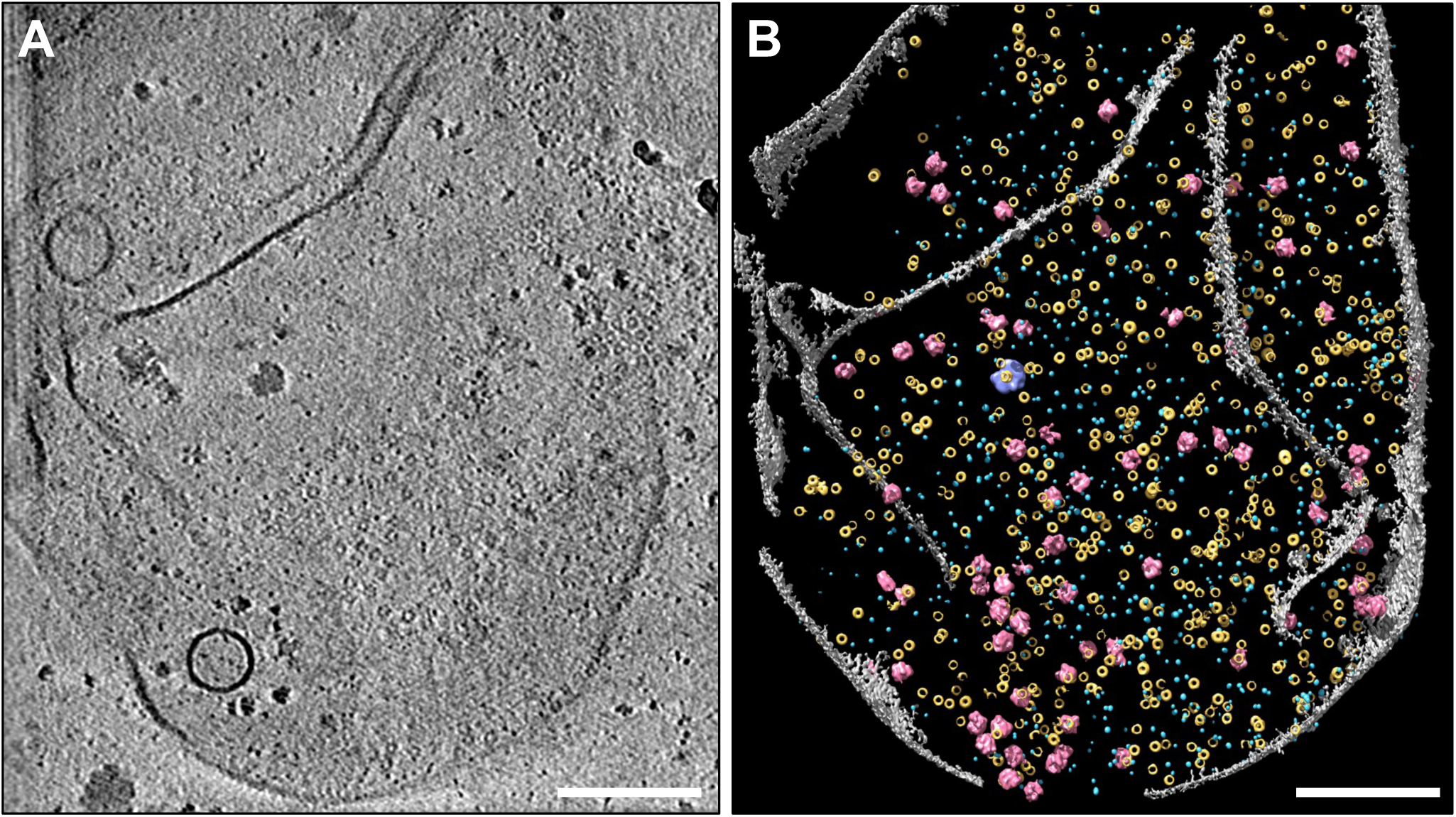
Annotation of a tomogram of fungal plasma membranes generated from caspofungin (CSF)-treated wild type spheroplasts. (**A**) Slice view of a plasma membr showing the distribution pattern of various molecular species. (**B**) Annotation and visualization of the drug-treated tomogram, depicting plasma membrane (gray), Pma1 hexamers (yellow), GS densities (light blue), ribosomes (pink) and glycogen granules (purple). Scale bars, 200 nm.

**Supplementary Figure 7.**
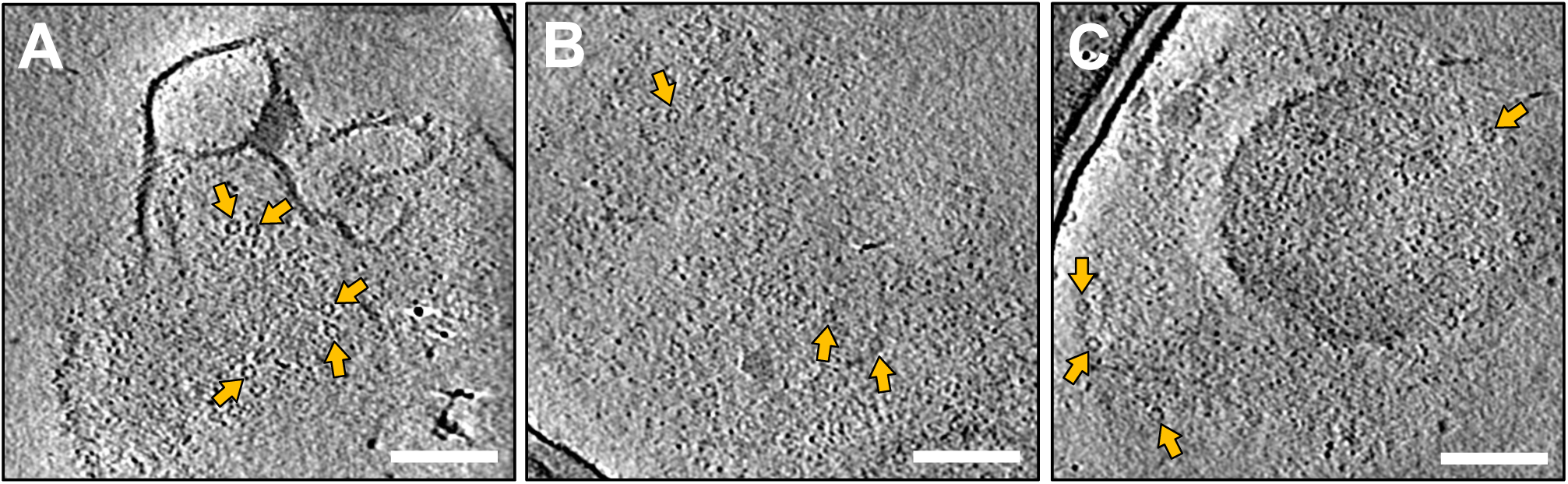
Cryo-ET of plasma membranes from Δfen1 mutant shows compromised membrane structures and dispersed Pma1 complexes. (**A-C**) Slice views of untreated plasma membranes from Δfen1 mutant cells. Loose Pma1 hexamers are indicated by yellow arrows. Scale bars, 100 nm.

## Supplementary Movies

**Supplementary Movie 1.** Slice views and annotation of a *C. glabrata* wild type plasma membrane tomogram showing the molecular sociology of fungal plasma membrane.

**Supplementary Movie 2.** Slice views of a *C. glabrata* cryo-lamellae tomogram revealing the cell wall, plasma membrane ultrastructure and subcellular features.

**Supplementary Movie 3.** Slice views and tomogram annotation of plasma membranes generated from *C. glabrata* wild type spheroplasts treated with caspofungin.

**Supplementary Movie S4.** Micropipette aspiration of untreated *C. glabrata* wild type spheroplasts.

**Supplementary Movie S5.** Micropipette aspiration of caspofungin-treated *C. glabrata* wild type spheroplasts.

## Materials and Methods

**Table 1.** *C. glabrata* strains used in this study.

| strain | parent | strain | plasmid | protein expressed | origin |
| --- | --- | --- | --- | --- | --- |
| CBS138 |  | wild type strain |  |  | ATCC |
| KH238 | CBS138 (ATCC2001) | 200989 $\Delta fks1$ + pCN-PDC1p-FKS1 | pCN-PDC1 | Fks1 overexpressing | Jiménez-Ortigosa et al., 2021 |
| IGCg1 | CBS138 (ATCC2001) | FKS1::NrsR-FKS1p-N-term_YFP-FKS1 | N/A | YFP-Fks1 expressing | this study |
| MVKCg1 | CBS138 (ATCC2001) | FKS1::NrsR-CgPDC1p*_N-term YFP-FKS1 | N/A | YFP-Fks1-overexpressing | this study |
| MVKCg9 | CBS138 (ATCC2001) | $\Delta FEN1::HygR$ | N/A | fen1 deletant mutant | this study |
| YHS26 | DPL1021 (ATCC90030) | $\Delta FEN1::HygR$ | N/A | fen1 deletant mutant | this study |

### Cell Culture Conditions and Preparation of Spheroplast Plasma Membranes

Cultures of *Candida glabrata* cells were grown in YPD (1% yeast extract, 2% peptone, 2% dextrose) or YPD supplemented with 100 µg/mL nourseothricin sulfate (Research Products International) (KH238, plasmid-carrying strain) to mid-logarithmic phase (OD_600_ = 2–3). The cells were harvested by centrifugation (5000*g* for 5 min), washed twice with water and incubated with 1% ß-mercaptoethanol for 1 h at 30°C with gentle shaking (Fisher Scientific). After incubation, the cells were washed twice with water by centrifugation and resuspended in buffer S (1M sorbitol, 10mM HEPES-NaOH, pH 6.5) in ratio of 4 ml of buffer/1 g wet cells. To remove the cell wall, the cell suspension was digested with lysing enzymes from *Trichoderma harzianum* (Sigma-Aldrich) and Zymolyase 20T (Zymo Research) at 100 mg and 100 units per 1 wet g of cells, respectively. The sample was then incubated overnight (14–16 hours) at room temperature with gentle shaking. The quality of spheroplasts generated was evaluated under a conventional light microscope. Spheroplasts were then collected by centrifugation at 3500*g* at 4°C and washed twice with buffer S, supplemented with an EDTA-free protease and phosphatase inhibitor cocktail (Roche) with 1 tablet per 10 ml buffer preparations.

To generate crude plasma membrane extracts, the spheroplast pellet was then resuspended in PBS, containing protease inhibitor and phosphatase inhibitor cocktails, for cell disruption. Cell lysis was confirmed by light microscopy. Centrifugation at 20,000*g* for 15 min at 4°C was performed to remove the presence of intact cells, spheroplasts, cellular debris, and other microsomal membranes. The crude membranes were resuspended in PBS supplemented with protease inhibitor and phosphatase inhibitor cocktails and used for subsequent mass spectrometry analysis and cryo-ET structural studies.

### Mass Spectrometry and Data Analysis

Crude membranes were prepared from wild type (CBS138) *C. glabrata* cells as described previously. The extracted membranes were run into an SDS-PAGE. Each gel band was subjected to reduction with 10 mM DTT for 30 min and then subjected to 60°C alkylation with 20 mM iodoacetamide for 45 min at room temperature. The sample was kept in the dark and further digested with trypsin (sequencing grade, Thermo Fisher Scientific) and incubated overnight at 37°C. The digested peptides were extracted twice with 5% formic acid, 60% acetonitrile and dried under vacuum.

The resulting peptides were analyzed by LC-MS using Nano LC-MS/MS (Dionex Ultimate 3000 RLSCnano System, Thermo Fisher Scientific) interfaced with Eclipse (Thermo Fisher Scientific). Samples were loaded onto a fused silica trap column Acclaim PepMap 100, 75 μm × 2 cm (Thermo Fisher Scientific) and washed with 0.1% trifluoracetic acid for 5 min with a flow rate of 5 µl/min. The trap column was brought in-line with an analytical column (nanoEase MZ peptide BEH C18, 130A, 1.7μm, 75 μm x 250 mm, Waters) for LC-MS/MS. Peptides were fractionated with a flow rate of 300 nl/min using a multistep linear gradient (4–15% Buffer A for 30 min [Buffer A: 0.2% formic acid], then 15–25% Buffer B for 40 min [Buffer B: 0.16% formic acid, 80% acetonitrile], followed by 25–50% Buffer B for 44 min, and 50–90% B in 11 min. For the following run, 4% Buffer B was used for 5 min.

The DIA (Data Independent acquisition) workflow was used for analysis of eluted peptides. The MS scan was set to a resolution of 120,000, with a scan range of 400–1200 and the AGC set to 3E6. An 8 m/z window was used to sequentially isolate (AGC of 4E5 and ion time set to auto) and fragment ions in the C-trap with a relative collision energy of 30. The fragments were recorded with resolution of 30,000. Raw data were analyzed using the predicted library of the *Candida glabrata* proteome provided on the Uniprot^52^ and the FASTA databases for library-free search using DIA NN 1.8.1 with recommended settings^53^. The results were filtered for both PEP (an estimate of the posterior error probability for the precursor identification, based on scoring with neural networks) filter <0.01 and PG.Q (Protein Group Q Value) filter <0.01.

The Uniprot *Candida glabrata* proteome database was used to determine the predicted total protein count from the *Candida glabrata* genome. To compile a list of only the most abundant plasma membrane proteins, we used subcellular localization information available in the Uniprot Knowledgebase^52^, the *Candida* Genome Database (CGD)^54^ and the *Saccharomyces* Genome Database (SGD)^55^ to generate a list of the most abundant fungal plasma membrane proteins present in our sample preparation. These protein candidates are either integral membrane proteins embedded in the cell membrane or attached to the plasma membrane by a lipid anchor. The multimeric state of the membrane protein candidates was determined by conducting literature review and searches for experimentally derived structures of the protein of interest within the RCSB Protein Data Bank (PDB)^56^. For protein candidates with no oligomerization information, we considered them as monomers. To account for the protein length (i.e., number of amino acids) and the oligomeric state, we normalized the protein group MaxLFQ values of the individual proteins to the molecular weight of their functional complex.

### Antifungal Susceptibility Testing and Caspofungin Treatment

Caspofungin (CSF) acetate (Selleck Chemical) was reconstituted in sterile ultrapure water (18.2 MΩ resistance) to yield a stock concentration of 1 mM. The stock was diluted as necessary for antifungal susceptibility tests.

Drug susceptibility microdilution assays were performed according to EUCAST guidelines with modifications^57,58^. Cells were grown and tested in RPMI 1640 media (Life Technologies) containing 2% of glucose without sodium bicarbonate, buffered with 165 mM MOPS–HCl to pH 7.0 at room temperature and filter sterilized. Tested drugs were serially diluted by 100 µl of 2× RPMI in a range of 2× concentrations in flat-bottom 96-well plates (Sarstedt). *C. glabrata* cultures at logarithmic growth phase (OD_600_ = 1–6) were diluted in ultrapure water to OD_600_ = 0.001 and distributed to the drug-containing plates by 100 µl (∼1500 CFU/well). Plates were incubated in moisture-controlling compartment at 37°C with shaking (150 RPM) in New Brunswick Innova® 42 incubator (Eppendorf) for 24 h (40 hours for slow growing strains). Plates were scanned at 600 nm using VERSAmax microplate reader (Molecular Devices). The MICs for CSF were defined as the concentration in the well with OD_600_ <5% above of the optical density for next (doubled) concentration point. Assays were performed in technical duplicates and two or more biological replicates.

We established the following CSF treatment conditions for each strain: 1 µg/ml for CBS138 and IGCg1, 2 µg/ml for KH238, and 6 µg/ml for MVKCg9. These concentrations are physiologically relevant and enable detectable changes for subsequent bioimaging studies. For live cell imaging, CBS138 and IGCg1 cells were first diluted to OD600 of 0.1 prior to treatment with 1 μg/ml CSF. Drug-treated intact cells were incubated at 200 RPM for 1 h at 37°C, washed with ultrapure water by centrifugation, and resuspended in PBS prior to live cell imaging. To investigate the effects of CSF on the spatial distribution of plasma membrane proteins *in situ*, we treated intact spheroplasts with CSF accordingly and incubated for 1 h at room temperature with gentle shaking. Following CSF treatment, crude plasma membranes were generated as described above.

### Vitrification of Crude Membranes and Intact Yeast Cells

Untreated and treated spheroplast membranes obtained from wild type (CBS138), Fks1-overexpressing (KH238), and the sphingolipid-deficient (MVKCg9) strains were mixed with 6 nm BSA gold tracers (Electron Microscopy Sciences) to facilitate tilt series alignment. An aliquot of 3.5 μl of the crude membrane sample was deposited onto freshly glow-discharged Quantifoil R2/2 200-mesh grids (Quantifoil Micro Tools GmbH,) using the Leica EM GP (Leica Microsystems) operated at 20°C and 95% humidity. Front blotting was performed for 4–6 sec prior to plunging the EM grid into liquid ethane.

For cryo-FIB experiments, intact MVKCg1 yeasts cells resuspended in PBS were vitrified on Quantifoil Au R2/2 London Finder 300-mesh grids (Quantifoil Micro Tools GmbH) following a similar vitrification procedure as described above for crude membranes. All plunge frozen grids were stored under liquid nitrogen until imaging.

### Cryo-FIB Lamellae Preparation

A modified protocol was used for micromachining vitrified *C. glabrata* cells ^59^. Grids were clipped using FIB Autogrids and loaded into an Aquilos 2 dual FIB/SEM instrument (Thermo Fisher Scientific). To improve sample conductivity, sample sputter coating was performed at 10 mA for 25 sec. In the microscope chamber, the grids were further deposited with organometallic Pt layers using the *in situ* GIS to provide a protective coating of the bulk sample. Following GIS deposition, the grids were sputter coated for an additional 7 sec at 10 mA to minimize charging artifacts.

Preliminary lamellae sites were identified using the MAPS software and AutoTEM Cryo was implemented to automate the milling process. A stepwise milling scheme was carried out as follows: rough milling steps of decreasing ion beam current (0.5 nA, 0.3 nA, 0.1 nA) was performed at all lamellae sites to remove bulk material above and below the designated lamella position. A final polishing of each lamella was carried out using 50 pA and 10 pA currents to target a nominal thickness of 200–300 nm. Grids containing yeast lamellae were stored under liquid nitrogen until further imaging.

### Cryo-ET and Tomogram Reconstruction

Micrographs and tilt series of the plasma membranes were collected on a Talos Arctica 200 kV microscope (Thermo Fisher Scientific) equipped with a post-column BioQuantum energy filter (slit width of 20 eV) and a K2 direct electron detector. SerialEM was used for automated data collection under the following conditions: 49,000× microscope magnification, spot size 8, 100-μm condenser and objective aperture, and with a nominal defocus range of −4–−6 μm^60^. The image pixel size was 2.73 Å/pixel. Tilt series ranged from −60° to +60° at 3° step increments. Tilt series were collected continuously from −60° to +60° with 3° step increments and in counting mode with a cumulative dose of 80–100 e^−^/Å^2^.

Tilt series of cryo-lamellae of intact *C. glabrata* cells were acquired on a Titan Krios 300 kV microscope (Thermo Fisher Scientific), operated a Selectris X energy filter (slit width set to 10 eV). SerialEM was used to perform automated data collection at a nominal magnification of 42,000× (pixel size of 3.041 Å/pixel) and defocus values ranging from −6–−7 μm. A dose-symmetric tilt series with 3° increments was used. Lamellae tomograms were recorded on a Falcon 4i direct electron detector with a cumulative dose of approximately 90 e^-^/Å^2^.

Raw movie frames were first motion-corrected with MotionCor2 software and tilt series stacks were generated^61^. Using EMAN tomography workflow, tilt series were aligned and reconstructed to generate 3D tomograms^62^. Tomograms were initially reconstructed with a binning factor of 4 to improve contrast.

### Tomogram Visualization and Annotation

Using the IMOD 3dmod interface, 2D slices were generated from 3D tomogram volumes^63^. For tomogram annotation, we used the EMAN2 convolutional neural network (CNN)-based protocol to annotate the plasma membrane, membrane protein structures and other subcellar components^23^. Using a high-contrast plasma membrane tomogram, we prepared different training datasets, i.e. positive and negative examples, to train a neural network for each structural feature. For Pma1, representative 2D patches of 24 positive examples and 132 negative examples were extracted and manually segmented to train the CNN. Due to the smaller size of putative GS densities, we selected 34 and 189 image tiles as positive and negative samples, respectively. We provided 10 positive and 109 negative examples for ribosomes, and 5 positive and 196 negative samples for glycogen granules. We used 64 × 64 pixel image tiles for training the CNN on Pma1, GS and ribosomes, and 112 × 112 pixel tiles for glycogen granules. The neural networks were trained for 20 iterations with a learning rate of 0.0001. We evaluated the training results prior to applying the trained neural networks to tomograms. If the training output appear suboptimal, we performed additional iterations of training until the accuracy of the CNN learning reached a satisfactory level. The trained neural networks were applied to annotate structural features from a CSF-treated plasma membrane tomogram.

To quantitatively assess CNN performance, we used manual annotation by an expert human annotator as ground truth. To visualize the manual annotation, we performed manual particle picking for each feature and mapped back the particles to the original coordinates in the tomogram. UCSF Chimera was used to visually compare the manual and CNN-based annotations at various isosurface thresholds^64^. True positives (TP), false negatives (FN) and false positives (FP) were determined and used to calculate the F1 score: TP/[TP + ½(FP + FN)].

### Statistical Analysis of Pma1 and Putative GS Density and Pma1 Clustering Behavior

To perform analysis of Pma1 and GS density on membranes, we used Pma1 and putative GS particles identified from our annotated tomogram. We first performed manual screening to remove false positive particles and incorporate missed particles. In this tomogram, we identified 254 Pma1 particles and 1369 GS particles. To quantify the membrane area of plasma membranes in our tomograms, we first generated a closed contour boundary model of the membrane using IMOD^63^. The density of Pma1 hexamers and putative GS monomers was then calculated as the particle count normalized to the area of the plasma membrane.

To characterize the clustering behavior of Pma1 hexamers in plasma membrane tomograms across different strains (CBS138 or KH238) and treatment conditions (with or without CSF treatment), we first manually boxed Pma1 particles in tomograms with sufficient image contrast and generated a list of 3D particle coordinates. We summarized the number of tomograms and particles boxed for each sample condition in **Supplementary Table 3**. We then performed nearest neighbor analysis of Pma1 hexamers in wild type plasma membranes. Within a Pma1 cluster, the nearest neighbor of a Pma1 hexamer was defined as the particle within the shortest linear distance. We manually estimated the number of distinct clusters present in the plasma membranes. For this calculation, we used information from the nearest neighbor analysis to define a cluster as four or more Pma1 hexamers in close proximity to one another with a distance of no more than 160–170 Å. We used Gaussian Mixture Model (GMM) and *k*-means clustering algorithms to evaluate our manual assignment of Pma1 clusters^65,66^. While GMM can capture the heterogeneity present in biological samples, *k*-means partitions the data into *k* distinct clusters and optimizes for intra-cluster variance. For GMM, we computed the Akaike Information Criterion (AIC) and Bayesian Information Criterion (BIC) for various clusters and evaluated which models had a lower score. For *k*-means clustering, we determined the optimal number of clusters by assessing the model’s inertia, which measures the compactness of the clusters. A lower inertia value suggests more distinct, densely packed clusters. Number of clusters determined from GMM-based and k-means were compared to our manual estimation (**Supplementary Table 2**).

To gain deeper insights into the clustering properties of Pma1 hexamer and compare distribution patterns across the four samples, we performed several statistical measures to assess the compactness of clusters: 1) intra-cluster distance and 2) cluster radius^67^. Given that the optimal cluster estimation from *k*-means clustering aligned better with our manual annotation, we calculated the average intra-cluster distance and cluster radius using *k*-means clustering and visualized the results as box plots. We applied the Kruskal-Wallis H test to determine whether these cluster metrics were significantly distinctive across the samples. For both average intra-cluster distance and cluster radius, the Kruskal-Wallis test yielded statistically significant differences with *p*-value < 0.05. To facilitate controlled comparisons and mitigate the potential for Type I errors, we proceeded with Dunn’s post-hoc test to pinpoint specific pairwise differences between the samples. We then performed power analysis through statistical simulation to ensure statistical validity of subsequent statistical tests. With 10,000 iterations, we simulated datasets based on the original sample’s means and standard deviations with each adjusted for an anticipated effect size of 0.8. This approach showed that the sample sizes were sufficient (estimated power of more than 0.7) to detect differences among the various groups, affirming the reliability of our statistical conclusions.

### Live Cell Fluorescent Imaging

*C. glabrata* cells from CBS138 and IGCg1 strains were cultured as described previously. Cells at mid-logarithmic growth were harvested and resuspended in PBS prior to application on coverslips for live cell fluorescence microscopy.

To reduce autofluorescence, 25 mm coverslips (0.17 ± 0.01 mm, Warner Instruments) were pre-cleaned as described previously with slight modifications^68^. Residual solvents were removed by drying using pressured N_2_. Next, pre-cleaned coverslips were plasma cleaned (Harrick Plasma Inc) for 10 minutes. Plasma-cleaned coverslips were placed into a clean container filled with a water chamber to maintain a humid environment. To immobilize the cells and minimize sample movement, coverslips were coated with 10 mg/ml concanavalin A (Thermo Fisher Scientific) for 5 min. Excess surfactant was gently removed and the coated coverslips were further dried with pressured N_2_. A volume of 200-500 μl of sample was applied onto the center region of the coverslips and incubated for 5 min to allow the cells to settle prior to visualization.

All microscopy experiments were conducted using a custom-built total internal reflection fluorescence (TIRF) microscope based on the Ti-E inverted microscope, with a high NA CFI-Apo 100X, NA 1.49 objective (Nikon) and an electron multiplying charge-coupled device (EMCCD) camera (iXon Ultra-888; Andor)^69^. The microscope was equipped with a 405-nm (OBIS 405 nm LX100 mW; Coherent) and a 488-nm (Genesis MX488-1000 STM; Coherent) lasers for the two-color fluorescence imaging required in this work. The microscope system has ∼40% light power delivery efficiency from the laser head to the sample. Differential interference contrast (DIC) imaging was conducted using a white light LED (LDB101F; Prior) and Nikon’s DIC modules. Multichannel images were obtained by triggered acquisition schemes, using acousto-optic tunable filter (AOTF) (AOTFnC-400.650-TN; Quanta-Tech), transistor-transistor logic (TTL) signal out of the EMCCD camera, a data acquisition card (PCIe-7852R; NI), and Nikon NIS-Elements software.

To stain for β-(1,3)-glucans in the cell wall, yeast cells incubated in 1.5 mg/ml aniline blue (Ward’s Science) for 5 min prior to imaging. Aniline blue was imaged using 405-nm laser excitation and a blue emission band-pass filter (ET475/m; Chroma), while YFP was imaged using 488-nm laser excitation and a yellow emission band-pass filter (ET530/30m; Chroma). Intact cells and spheroplasts were first inspected in the DIC channel and then switched to the fluorescence channel to optimize imaging parameters (e.g. laser power, illumination angle, camera exposure, etc.) for TIRF imaging. For DIC imaging, exposure time was typically set to 50 ms. For imaging aniline blue, the 405-nm laser was used at power settings of 10 mW, coupled with exposure time of 500 ms. For imaging YFP, the 488-nm laser was used at power settings of 10 mW, coupled with exposure of 500 ms. The electron multiplication (EM) gain of the EMCCD camera was set to 50 and 230 for DIC and fluorescence imaging, respectively. The Perfect Focus System (Nikon) was implemented to actively stabilize focus drift during image acquisition. Digital image analysis was performed using Fiji ImageJ^70,71^.

### Measurement of Surface Tension by Micropipette Aspiration (MPA)

Micropipettes were pulled from glass capillaries using a pipette puller (PUL-1000, World Precision Instruments (WPI)). The pipette tip was cut to an opening diameter smaller than the average spheroplast radius (∼2.5 µm) between 1.5–2 μm.

Micropipette aspiration (MPA) and imaging were performed on a Ti2-A inverted fluorescent microscope (Nikon) equipped with a motorized stage and two motorized micromanipulators (PatchPro-5000, Scientifica). Harvested *C. glabrata* spheroplasts were maintained in Buffer S, an osmotically stabilizing buffer, to prevent lysis. A micropipette was filled with the same buffer used for spheroplasts using a MICROFIL needle (WPI) and then mounted onto a micromanipulator connected to the pressure control (LU-FEZ-N069, Fluigent).

The zero pressure of the system was calibrated before each MPA experiment, using a dilute solution of small particles. The zero pressure (P0) was set according to the point when the particles undergo random Brownian motion in the micropipette. After calibration of the aspiration pressure, 100–200 μl sample of the spheroplast sample was loaded onto the center of a glass-bottom dish (ES56291, Azer Scientific) and MilliQ water was added to the edge of the dish to minimize sample evaporation. A calibrated micropipette was then moved to a spheroplast with a diameter of approximately 5 µm.

During aspiration measurements, sequential stepwise suction pressures were applied to deform the spheroplast. Aspiration pressure was gradually increased every 20–30 seconds. The surface tension σ was calculated as σ = ΔP · R_p_/ [2(1 - R_p_/R_c_)] where R_p_ and R_c_ are the micropipette and cell radius, respectively, and ΔP is the aspiration pressure at which the length of the spheroplast aspirated into the micropipette is closest to R_p_. Deformation of the spheroplast membrane was recorded using a 60X objective at Hz (ORCA-Flash 4.0, Hamamatsu) through transmitted light imaging. To capture the effect of CSF treatment on biophysical properties of the plasma membrane, spheroplasts were treated with 1 µg/ml CSF for 15 min prior to micropipette aspiration and imaging. ImageJ was used to track the shape parameters (L_p_, R_p_, R_c_) of the aspirated spheroplasts^70,71^.

